# Hecaton: reliably detecting copy number variation in plant genomes using short read sequencing data

**DOI:** 10.1101/720805

**Authors:** Raúl Wijfjes, Sandra Smit, Dick de Ridder

**Author notes:** Corresponding author: Dick de Ridder.

## Abstract

Copy number variation (CNV) is thought to actively contribute to adaptive evolution of plant species. While many computational algorithms are available to detect copy number variation from whole genome sequencing datasets, the typical complexity of plant data likely introduces false positive calls.

To enable reliable and comprehensive detection of CNV in plant genomes, we developed Hecaton, a novel computational workflow tailored to plants, that integrates calls from multiple state-of-the-art algorithms through a machine-learning approach. In this paper, we demonstrate that Hecaton outperforms current methods when applied to short read sequencing data of *A. thaliana*, rice, maize, and tomato. Moreover, it correctly detects dispersed duplications, a type of CNV commonly found in plant species, in contrast to several state-of-the-art tools that erroneously represent this type of CNV as overlapping deletions and tandem duplications. Finally, Hecaton scales well in terms of memory usage and running time when applied to short read datasets of domesticated and wild tomato accessions. Hecaton provides a robust method to detect CNV in plants. We expect it to be of immediate interest to both applied and fundamental research on the relationship between genotype and phenotype in plants.

## 1 Introduction

Phenotypic variation between individuals of the same plant species is caused by a host of different types of genetic variation, including single nucleotide polymorphisms (SNPs), small insertions and deletions, and larger structural variation. One major class of structural variation is copy number variation (CNV), which is defined as deletions, insertions, tandem duplications and dispersed duplications of at least 50 bp. CNV comprises a large part of the genetic variation found within plant populations and is thought to play a key role in adaptation and evolution (Zmienko et al., 2014). One clear example of such adaptive evolution is presented by the weed species *A. palmeri*, which rapidly became resistant to a widely used herbicide through amplification of the EPSPS gene, resulting in increased expression (Gaines et al., 2010). Similar relationships between CNV and adaptation were found in domesticated crop species (Gabur et al., 2019), indicating that CNV may offer a pool of genetic variation that can be used to improve crop cultivars.

Given the increasing interest of the plant research community in CNV (Zmienko et al., 2014; Gabur et al., 2019; Lye and Purugganan, 2019), the question arises whether current methods accurately detect copy number variants (CNVs) in plants. Currently, CNVs are mainly analyzed by whole genome sequencing (WGS). After a sample of interest has been sequenced and the resulting sequencing data has been aligned to a reference genome, computational methods can extract various signals from the alignments to detect CNV between the sample and the reference (Alkan et al., 2011). While long reads are better suited for detecting CNVs than short paired-end reads (Sedlazeck et al., 2018; De Coster et al., 2019), sequencing data of plants is still commonly generated using short read sequencing platforms, due to their far lower cost.

Although current state-of-the-art CNV detection algorithms generally perform well when applied to human datasets (Kosugi et al., 2019), the typical complexity of plant data likely introduces false positive calls. First, reference genome assemblies of plants generally contain a larger number of gaps than the human reference genome, as plant genomes are difficult to assemble due to their repetitive nature. Yet, the genomic sequence contained in such gaps is still present in WGS data of samples. The reads representing this sequence generally share high similarity with other assembled regions of the reference, to which they are incorrectly aligned as a result. Second, sampled plant genomes can differ significantly from reference genome assemblies, particularly if samples represent out-bred or natural accessions. If a region in a sample genome has undergone several mutations relative to the reference, reads sequenced from this region may map to a different region than the one it is syntenic to. This is particularly likely to happen if the region that the reads originated from is highly repetitive. Third, several CNV detection algorithms erroneously process alignments resulting from dispersed duplications (Zhao et al., 2016). We expect that this issue introduces a significant number of false positives when working with plant data, as duplication and transposition of genomic sequences is considered to be one of the main drivers of adaptive evolution in plants (Lisch, 2013).

To enable reliable and comprehensive detection of copy number variants in plant genomes, we developed Hecaton, a novel computational workflow that combines several existing detection methods, specifically tailored to detect CNV in plants. Combining methods generally results in higher recall and precision than using a single tool (Mills et al., 2011; Chaisson et al., 2019), as the recall and precision of individual tools varies among different types and sizes of CNVs, depending on their algorithmic design (Kosugi et al., 2019). However, determining the optimal strategy to integrate different methods is not straightforward. A suboptimal integration approach may yield only a small gain of precision, while significantly decreasing recall (Lee et al., 2018; Kosugi et al., 2019). Hecaton tackles this challenge in two ways. First, it makes use of a custom post-processing step to correct erroneously detected dispersed duplications, which are systematically mispredicted by some state-of-the-art tools. Second, it utilizes a machine-learning model which classifies detected calls as true and false positives by leveraging several features describing a detected CNV call, such as its type and size, along with concordance among the callers used to detect it. In this paper, we demonstrate that Hecaton outperforms existing individual and ensemble computational CNV detection methods when applied to plant data and provide an example of its utility to the plant research community.

## 2 Materials and methods

### 2.1 Selected CNV calling tools

To maximize the performance of Hecaton, we combine predictions of a diverse set of popular, open-source tools that complement each other in terms of the signals and strategies used to call CNVs. The selected tools include Delly (Rausch et al., 2012) (version 0.7.8), GRIDSS (Cameron et al., 2017) (version 1.8.1), LUMPY (Layer et al., 2014) (version 0.2.13), and Manta (Chen et al., 2015) (version 1.4.0). Delly detects CNVs using discordantly aligned read pairs and refines the breakends of detected events using split reads. LUMPY improves upon this method by integrating both of these signals to detect CNVs, as opposed to using them sequentially. Manta and GRIDSS further enhance this strategy by performing local assembly of sequences flanking breakends identified by discordantly aligned read pairs and split reads. We considered including CNVnator (Abyzov et al., 2011) (version 0.3.2), Control-FREEC (Boeva et al., 2011) (version 10.4), and Pindel (Ye et al., 2009) (version 0.2.5b9). Pindel was dropped after showing an excessively long run time when applied to simulated high coverage datasets. CNVnator and Control-FREEC were excluded as they performed poorly during evaluations (Additional File 1: Figure S1).

### 2.2 Implementation of Hecaton

Hecaton is a workflow specifically designed to reliably detect CNVs in plant genomes. We aimed to implement it in such a manner that it is both reproducible and easy-to-use. To this end, Hecaton is run with a single command using the Nextflow (Di Tommaso et al., 2017) framework, which provides a unified method to chain together and parallelize the different processes that are executed. It consists of three stages: calling, post-processing, and filtering (Figure 1). Currently, Hecaton only supports the four CNV detection algorithms used during the calling stage, but can be relatively easily extended to include other tools.

**Figure 1:**
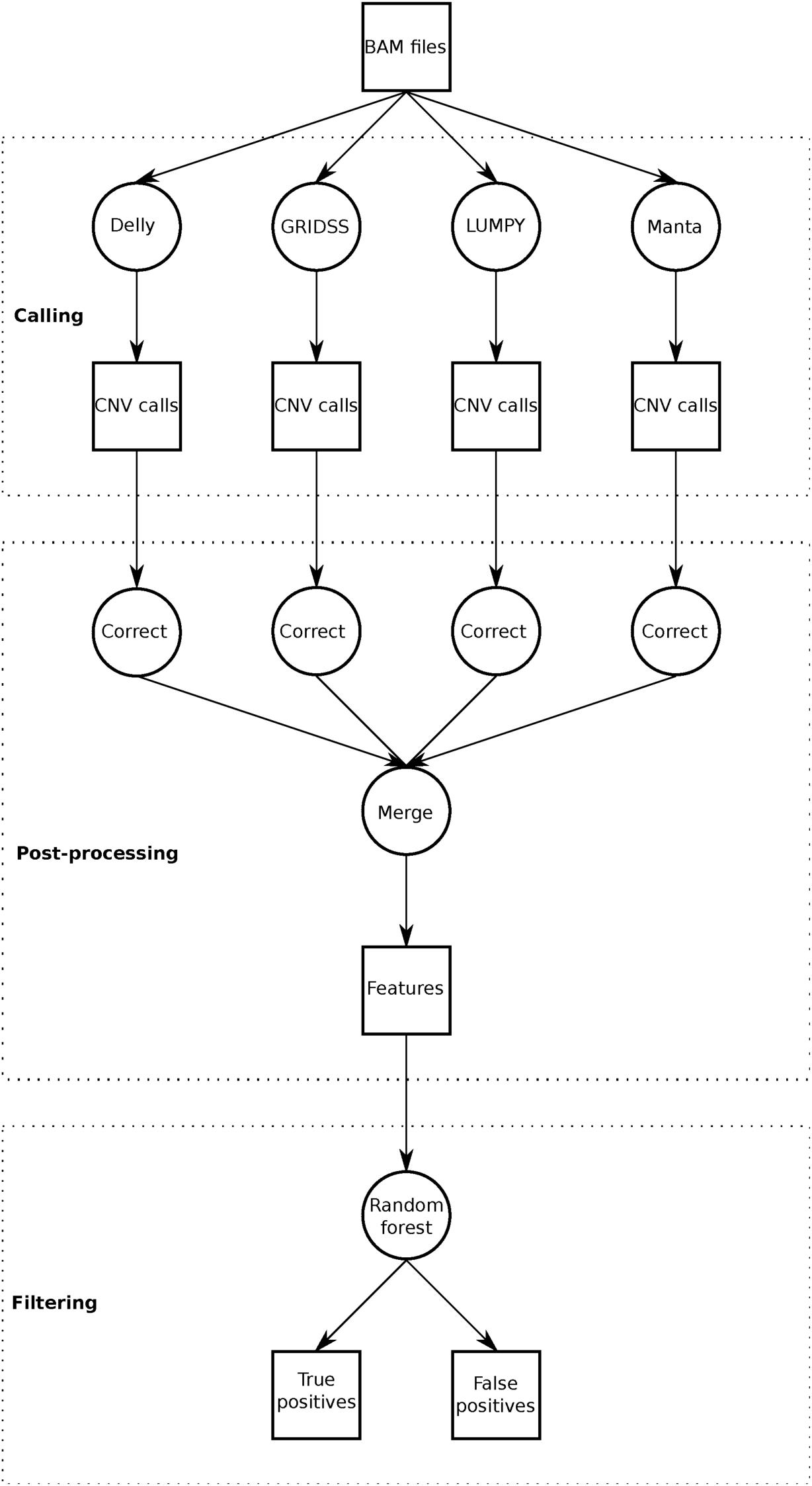
Overview of Hecaton. CNVs are first called using four different tools. The resulting calls are corrected and merged into a set of features. These features are used by the random forest model to discriminate between true and false positives.

#### 2.2.1 Stage 1: Calling

The calling stage takes paired-end Illumina WGS data of a sample of interest and a reference genome as input and calls CNVs between the sample and reference using four different tools. First, it aligns the sequencing data to the reference using the Speedseq pipeline (Chiang et al., 2015) (version 0.1.2) with default parameters. This pipeline utilizes bwa mem (Li, 2013) (version 0.7.10-r789) to align reads, SAMBLASTER (Faust and Hall, 2014) (version 0.1.22) to mark duplicates and Sambamba (Tarasov et al., 2015) (version 0.5.9) to sort and index BAM files. The resulting sorted BAM file is processed by Delly, LUMPY, Manta and GRIDSS to call CNVs. Each of these tools is run with default parameters, except for the number of supporting reads required by LUMPY and Manta for a CNV to be included in the output (lowered to 1 to maximize sensitivity). Delly and GRIDSS do not apply any filters by default. The final output of the calling stage consists of four VCF files containing CNVs, one for each tool.

#### 2.2.2 Stage 2: Post-processing

The post-processing stage of Hecaton serves three purposes. First, it provides an automated method to process the output files of different tools using a common representation, which is necessary to properly integrate them. Second, it corrects dispersed duplications that have been detected by CNV tools as overlapping deletions and tandem duplications by mistake. Third, it merges calls produced by different tools that likely correspond to the same CNV event.

The common representation of CNVs used by Hecaton is based on the concept that each structural variant can be represented as a set of novel adjacencies. A novel adjacency is defined as a pair of bases that are adjacent to each other in the genome of a sample of interest, but not in the genome of the reference to which the sample is compared. Bases that are linked by a novel adjacency are called breakends and two breakends that corresponds to the same adjacency are referred to as mates. Although Delly, GRIDSS, LUMPY, and Manta all generate a VCF file as output, the way in which CNV calls and the evidence supporting them are represented in this file is different for each tool. For example, the output of Delly, LUMPY, and Manta contains both CNVs and breakends, while that of GRIDSS solely consists of breakends that need to be converted to CNVs by the user.

To convert the output of each tool to a common CNV format and correct erroneous dispersed duplications, Hecaton reclassifies the adjacencies underlying the CNV calls produced by each tool. First, it infers and collects adjacencies from all sets of CNVs generated during the calling stage. For example, it represents deletions as a single adjacency containing two breakends positioned on the 5’ and 3’ end of the deleted sequence. Next, it clusters adjacencies generated by the same tool of which the breakpoints are located within 10 bp of each other on either the 5’ end or 3’ end, as these are likely to be part of the same variant. Finally, it converts each cluster to a deletion, insertion, tandem duplication, or dispersed duplication, based on the relative positions of the breakends and the orientation of the sequences that are joined in a cluster. Deletions, insertions, and tandem duplications are represented by single adjacencies, while dispersed duplications are represented by two (Additional File 1: Figure S2). As the objective of Hecaton is to detect CNV and not any other form of structural variation, it excludes any set of adjacencies that cannot be classified as one of these four types from further analysis. However, Hecaton can be extended to support additional types of structural variation if needed.

Hecaton collapses calls produced by different tools that are likely to correspond to the same CNV event. Calls are merged if they fulfill all of the following conditions: they are of the same type; their breakpoints are located within 1000 bp of each other on both the 5’ and 3’ end; they share at least 50% reciprocal overlap with each other (does not apply to insertions); and the distance between the insertion sites is no more than 10 bp (only applies to dispersed duplications and insertions). The regions of the merged calls are defined as the union of the regions of the “donor” calls. For instance, one call that covers positions 12-30 and one call that covers positions 14-32 are merged into a call covering positions 12-32. The number of discordantly aligned read pairs and split reads supporting a merged call are both defined as the median of the numbers of the donor calls. The final result of the post-processing stage is a single BEDPE file containing all merged calls.

#### 2.2.3 Stage 3: Filtering

In the filtering stage, Hecaton applies a machine-learning model to remove erroneous CNV calls. First, it generates a feature matrix that represents the set of merged calls. The rows of the matrix correspond to CNV calls and the columns correspond to features (Additional file 2: Table S1), which are extracted from the INFO and FORMAT fields of the VCF file containing the calls.

Hecaton classifies calls as true or false positives using a random forest model. We chose to implement this particular type of machine-learning model, as it outperformed a logistic regression model and a support vector machine. The model assigns a probabilistic score to each merged call based on the set of features defined for it. These scores are posterior probability estimates of calls being true positives and range between 0 and 1. Calls with scores below a specified user-defined cutoff are dropped, producing a BEDPE file containing the final output of Hecaton.

To obtain a random forest model that strikes a good balance between sensitivity and precision, we trained it using a set of CNVs detected from real WGS data for which the labels (true or false positive) were known, based on long read data (see Additional file 3: Supplementary Methods for details on the validation procedure). We did not include CNVs obtained from simulated data in the ground truth set, as the sensitivity and precision attained by Delly, LUMPY, Manta, and GRIDSS on such data generally does not accurately reflect their performance in real scenarios. For example, LUMPY and Manta obtained almost perfect precision when we applied them to simulated datasets with minimum filtering, if dispersed duplications were excluded from the simulation. They showed significantly lower precision in previous benchmarks when applied to real human data (Layer et al., 2014; Chen et al., 2015).

The training and testing set were constructed by running the calling and post-processing stages of Hecaton on Illumina data of an *A. thaliana* Col-0–Cvi-0 F1 hybrid and a sample of the *Japonica* rice Suijing18 cultivar (Additional file 2: Table S2). We detected CNVs in these samples relative to the *A. thaliana* Col-0 (version TAIR10) and *O. sativa Japonica* (version IRGSP-1.0) reference genome. As we aimed to maximize the performance of the model for low coverage datasets in particular, we subsampled these datasets to 10x coverage using seqtk (Li, 2012). Calls were labeled as true or false positives using long read data of the same samples (See Additional file 3: Supplementary Methods for details). To obtain a test set, we held out calls located on chromosomes 2 and 4 of *A. thaliana* and chromosomes 6, 10, and 12 of *O. sativa*, using the remaining calls as the training set. In order to obtain a model that generalizes to multiple plant species, one single model was trained using both Col-0–Cvi-0 and Suijing18 calls. The training set contained 4983 deletions, 393 insertions, 604 tandem duplications and 106 dispersed duplications, while the test set contained 2291 deletions, 174 insertions, 292 tandem duplications and 44 dispersed duplications.

We implemented the random forest model in Python using the scikit-learn package (Pedregosa et al., 2011) (version 0.19.1). The hyperparameters of the model (*n_estimators, max_depth*, and *max_features*) were selected by doing a grid search with 10-fold cross-validation on the training set, using the accuracy of the model on the validation data as optimization criterion.

### 2.3 Benchmarking

The performance of Hecaton was compared to that of current state-of-the-art tools using short read data simulated from rearranged versions of the *S. lycopersicum* Heinz 1706 reference genome of tomato (The Tomato Genome Consortium, 2012); the testing set constructed from *A. thaliana* Col-0–Cvi-0 and rice Suijing18; and real short read data of *A. thaliana* L*er*, maize B73, and several tomato samples (Additional file 2: Table S2). We determined the sensitivity and precision of tools with two validation methods that use long read data: VaPoR (Zhao et al., 2017) and Sniffles (Sedlazeck et al., 2018). See Additional file 3: Supplementary Methods for full details.

## 3 Results and Discussion

We present Hecaton, a novel computational workflow to reliable detect CNVs in plant genomes (Figure 1). It consists of three stages. In the first stage, it aligns short read WGS data to a reference genome of choice and calls CNVs from the resulting alignments using Delly, GRIDSS, LUMPY, and Manta, four state-of-the-art tools that complement each other in terms of their methodological set-up. In the second stage, Hecaton corrects dispersed duplications that are erroneously represented by these tools as overlapping deletions and tandem duplications. In the final stage, Hecaton filters calls by using a random forest model trained on CNV calls validated by long read data. Below, we first describe how the design of Hecaton allows it to outperform the current state-of-the-art and then we will present an application of Hecaton to crop data.

### 3.1 Hecaton accurately detects dispersed duplications

Dispersed duplications are defined as duplications in which the duplicated copy is found at a genomic region that is not adjacent to the original template sequence. Such variants are frequently found in plants, as plant genomes typically contain a large number of class I transposable elements that propagate themselves through a “copy and paste” mechanism. While dispersed duplications may play an important role in the adaptive evolution of plants (Lisch, 2013), they can also introduce a significant number of false positives, if they are not taken into account while calling CNVs. To show the impact of this problem, we applied Delly, GRIDSS, LUMPY, and Manta to short read data simulated from modified versions of the *S. lycopersicum* Heinz 1706 reference genome containing different types of CNVs at known locations.

As Delly, LUMPY, and Manta systematically mispredict dispersed duplications, they attained low precision when applied to simulated data (Figure 2a). We hypothesize that these tools misinterpret the complex patterns of signals resulting from intrachromosomal dispersed duplications during alignment (Additional file 1: Figure S2), as the false positives mostly corresponded to overlapping pairs of large deletions and tandem duplications (Figure 2b) that cover the sequence located between the template sequence and insertion sites of simulated intrachromosomal dispersed duplications. Such signals consist of novel adjacencies, pairs of bases that are adjacent to each other in the genome of the sample of interest, but not in the genome of the reference to which the sample is compared. Deletions, insertions, and tandem duplications generate a single novel adjacency as a signal. Dispersed duplications, however, generate two novel adjacencies. Delly, LUMPY, and Manta likely process these adjacencies in isolation, resulting in overlapping deletion and tandem duplication calls.

**Figure 2:**
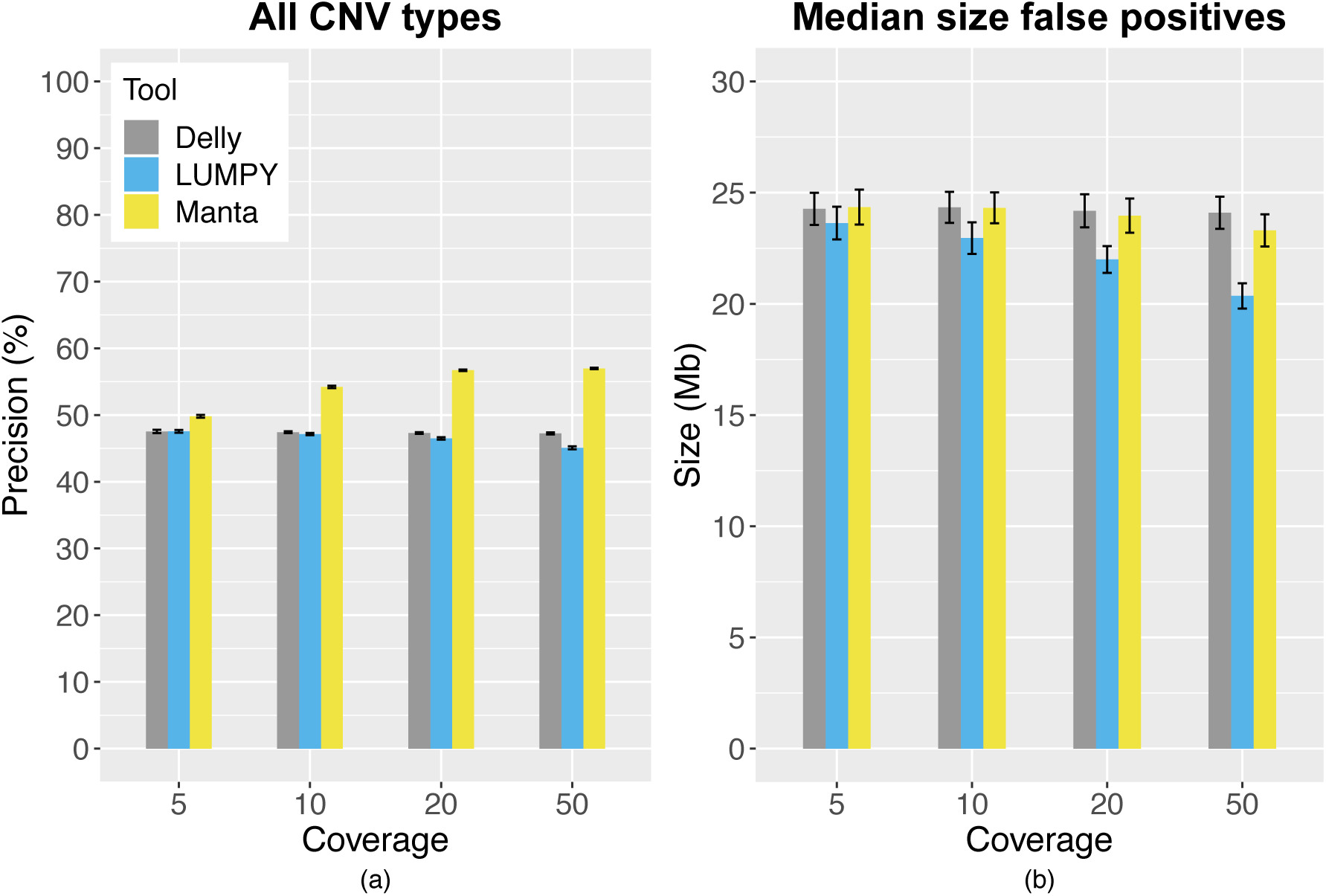
Performance of Delly, LUMPY, Manta, and GRIDSS on data simulated from diploid rearranged tomato genomes. Performance metrics are reported as the mean over all 10 simulations with error bars depicting the standard error of the mean. The overall precision of Delly, LUMPY, and Manta was low (a) and false positives generally consisted of large CNVs having a size of several tens of Mbs (b). These corresponded to pairs of large deletions and tandem duplications that covered the sequence located between the template sequence and insertion sites of intrachromosomal dispersed duplications.

The post-processing step of Hecaton corrects dispersed duplications that are erroneously predicted by Delly, LUMPY, and Manta, which significantly improves their performance. It recovered both intrachromosomal and interchromosomal dispersed duplications when applied to simulated data (Figure 3a). Moreover, as the post-processing step replaces false positive deletions and tandem duplications by true positive dispersed duplications, it strongly increases the precision of Delly, LUMPY, and Manta (Figure 3b). The post-processing step also correctly predicts dispersed duplications from the output of GRIDSS, which does not yield CNVs as output, but the adjacencies underlying them (Figure 3). Post-processing the adjacencies reported by GRIDSS in isolation resulted in a similar trend as seen for Delly, LUMPY, and Manta, underlining the importance of correctly interpreting the signals generated by dispersed duplications.

**Figure 3:**
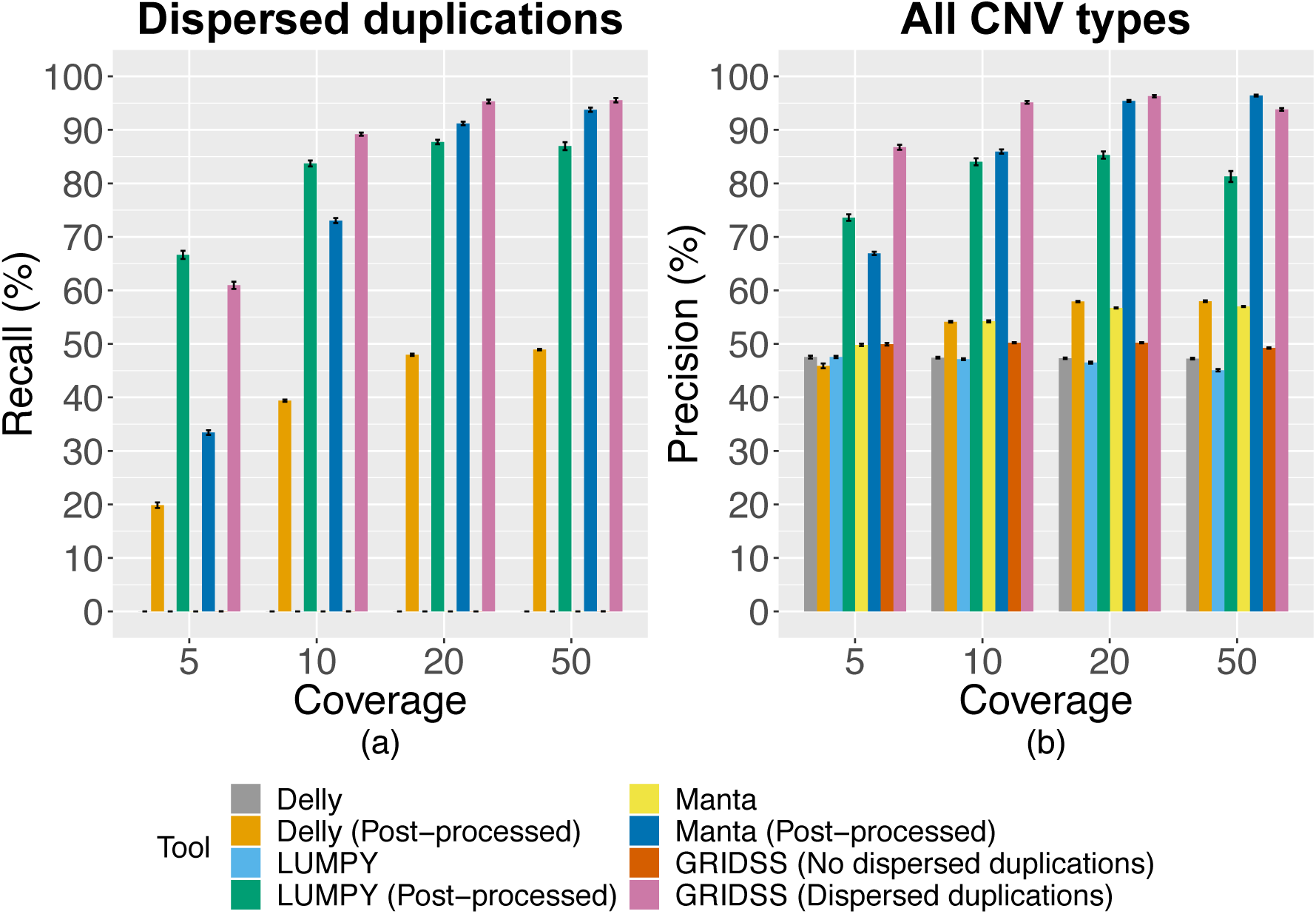
Performance of the post-processing step of Hecaton on data simulated from diploid rearranged tomato genomes. Performance metrics are reported as the mean over all 10 simulations with error bars depicting the standard error of the mean. Results of GRIDSS were generated by processing adjacencies in isolation (no dispersed duplications) or by processing them in clusters (dispersed duplications). (a) Recall of CNV calling tools for dispersed duplications, before and after post-processing. The post-processing script of Hecaton recalled dispersed duplications not originally found in the output of Delly, LUMPY, Manta. (b) Overall precision of CNV calling tools, before and after post-processing. The post-processing stage of Hecaton significantly increased the precision of tools by replacing pairs of overlapping false positive deletions and tandem duplications by true positive intrachromosomal dispersed duplications.

The performance of the post-processing step improved with coverage (Figure 3), as it fails to detect dispersed duplications if one or both of the adjacencies resulting from them are missing from the output of Delly, LUMPY, Manta, or GRIDSS. In line with this observation, the post-processing script detected a lower number of dispersed duplications simulated at low allele dosage compared to those simulated at high dosage (Additional file 1: Figure S3), as the effective coverage of variant alleles decreases when they are present in few haplotypes. If only one of the two adjacencies could be detected, the post-processing script classified it as a false positive deletion, false positive tandem duplication, or generic breakend.

### 3.2 Hecaton generally outperforms state-of-the-art CNV detection tools

Intuitively, it makes sense to combine the output of multiple CNV detection tools, as they typically generate complementary results when applied to the same dataset (Lin et al., 2014). However, designing a method that optimally integrates tools is not trivial. In a past benchmark, an ensemble strategy that combined tools through a majority vote did not significantly improve upon the best performing individual tool (Lee et al., 2018). Here, we demonstrate the benefits of using a machine-learning approach, which aggregates and filters calls based on features including size, type and level of support from different tools. We trained machine-learning models using CNVs detected from 10x coverage short read data of a highly heterozygous *A. thaliana* Col-0–Cvi-0 sample and a Suijing18 rice sample. The labels (true or false positive) of these CNVs were determined using long read data of the same samples. This approach generated accurate validations of calls detected from the simulated *S. lycopersicum* Heinz 1706 datasets.

The machine-learning approach used during the filtering stage of Hecaton integrates calls of Delly, LUMPY, Manta, and GRIDSS in such a manner so that it outperforms each individual tool. When applied to *A. thaliana* Col-0–Cvi-0 and Suijing18 rice calls detected on chromosomes that were held out from model training, it generally attained a more favourable combination of sensitivity and precision across a broad spectrum of thresholds and different CNV types (Figure 4). For example, at a precision level of 80%, Hecaton detected 43 true positive tandem duplications, while the best performing state-of-the-art tool, GRIDSS, detected only 19. Our results agree with previous work in which a method that carefully merges calls of different CNV calling tools attained a higher precision and sensitivity than any of the individual tools (Mills et al., 2011). As the approach performed about equally well when using a random forest model trained on either 10x or 50x coverage data (Additional file 1: Figure S4), the random forest framework itself is the main driver of the improvement, rather than the sequencing coverage used to train the models. To check whether the improved performance held more generally, we applied Hecaton to an Illumina dataset of *A. thaliana* L*er*, a sample that was completely independent from model training. It again improved upon the performance of individual tools (Additional file 1: Figure S5), corroborating the results observed in *A. thaliana* Col-0–Cvi-0 and Suijing18 rice.

**Figure 4:**
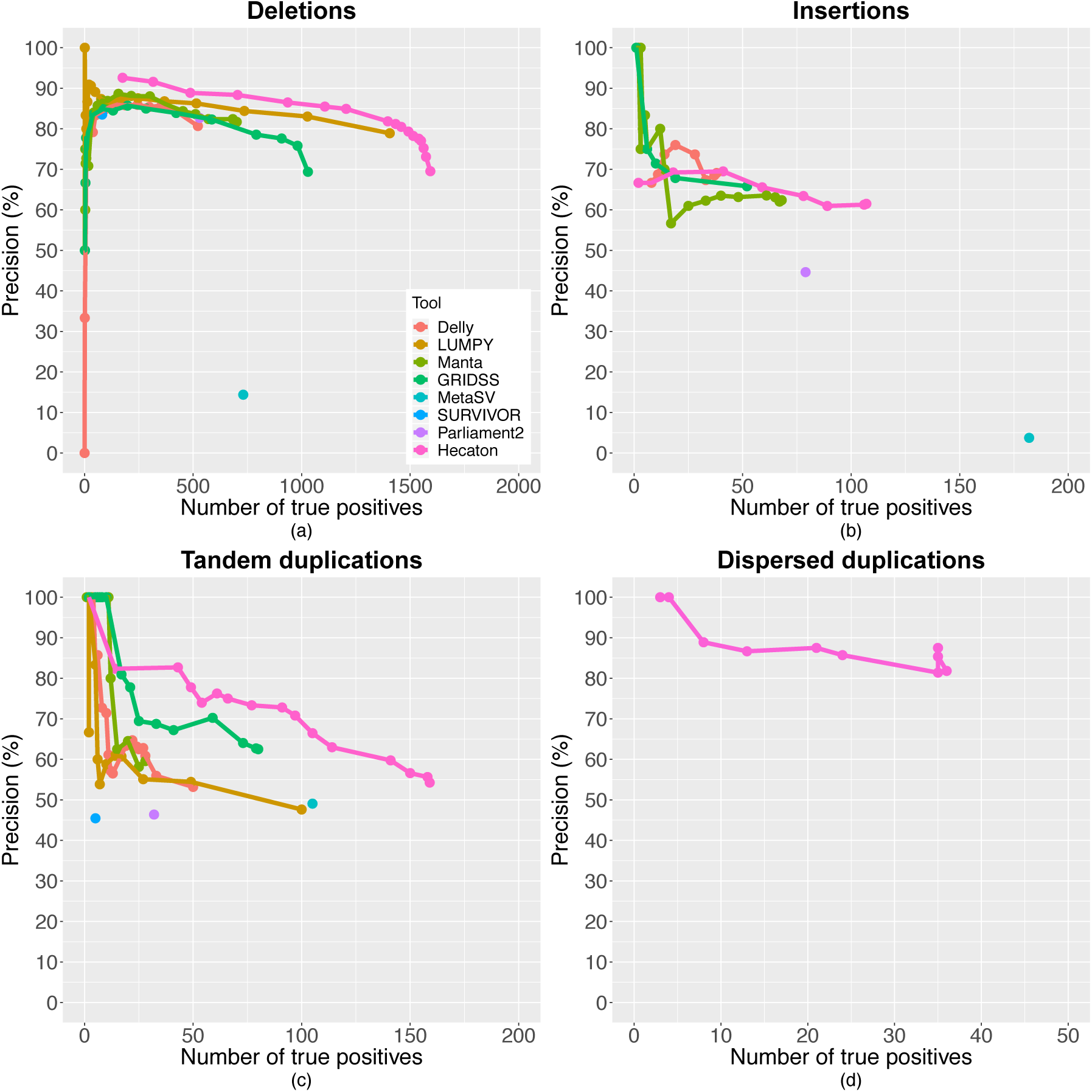
Performance of different CNV detection algorithms on the test set containing Col-0–Cvi-0 and Suijing18 CNV events. Precision-recall curves of Delly, LUMPY, Manta, and GRIDSS were constructed by varying the minimum number of discordantly aligned read pairs and/or split reads supporting each call in the set. The curve of Hecaton was produced by varying the threshold of the probabilistic score used to define calls as true positives. Performance is shown separately for deletions (a), insertions (b), tandem duplications (c), and dispersed duplications (d). Curves of LUMPY and SURVIVOR are missing for insertions, as these tools are unable to detect this type of CNV. Curves are missing for all tools besides Hecaton for dispersed duplications, as Hecaton is the only tool that can detect this type of CNV, owing to its post-processing stage. Hecaton generally improved recall and precision compared to Delly, LUMPY, Manta, and GRIDSS. Moreover, it significantly outperformed MetaSV, SURVIVOR, and Parliament2, three ensemble approaches applicable to plant data.

Besides outperforming individual tools, the machine-learning approach employed by Hecaton significantly improved upon current state-of-the-art ensemble methods that are applicable to, but not specifically designed for plant data. It attained a better combination of precision and sensitivity than MetaSV (Mohiyuddin et al., 2015), SURVIVOR (Jeffares et al., 2017), and Parliament2 (Zarate et al., 2018), three alternative approaches that aggregate the results of different CNV detection tools, when applied to datasets of Col-0–Cvi-0 and Suijing18 (Figure 4). The poor performance of MetaSV and SURVIVOR sharply contrasts with the good performance they showed in the benchmarks of the publications describing them (Mohiyuddin et al., 2015; Jeffares et al., 2017). One possible reason for this discrepancy could be that both tools were evaluated in these benchmarks using simulated data, which likely does not accurately reflect the distribution of CNVs in real data.

To evaluate Hecaton on more distantly related and repetitive genomes than those of *A. thaliana* and rice, we used it to detect CNVs between the two maize accessions Mo17 and B73. As a large fraction of calls could not be validated using long read data, due to the highly repetitive nature of the Mo17 assembly (Additional File 2: Table S3), we only report performance metrics for calls that overlap for at least 50% of their length with genes or the 5000 bp interval upstream or downstream of genes. We believe that this subset of calls still yields a representative measure of performance, as downstream analysis of CNVs detected by short reads generally focuses on genic, non-repetitive regions. Consistent with the results of our previous benchmarks, Hecaton attained a better combination of sensitivity and precision compared to both individual state-of-the art tools and ensemble approaches (Figure 5). For example, at a precision level of 90%, it detected a higher number of true positive deletions (13991) than LUMPY (11190), the second-most sensitive approach for deletions at that level of precision. The large number of CNVs detected by Hecaton between Mo17 and B73 confirms the extensive structural variation between the two accessions found by a whole genome alignment based approach (Sun et al., 2018).

**Figure 5:**
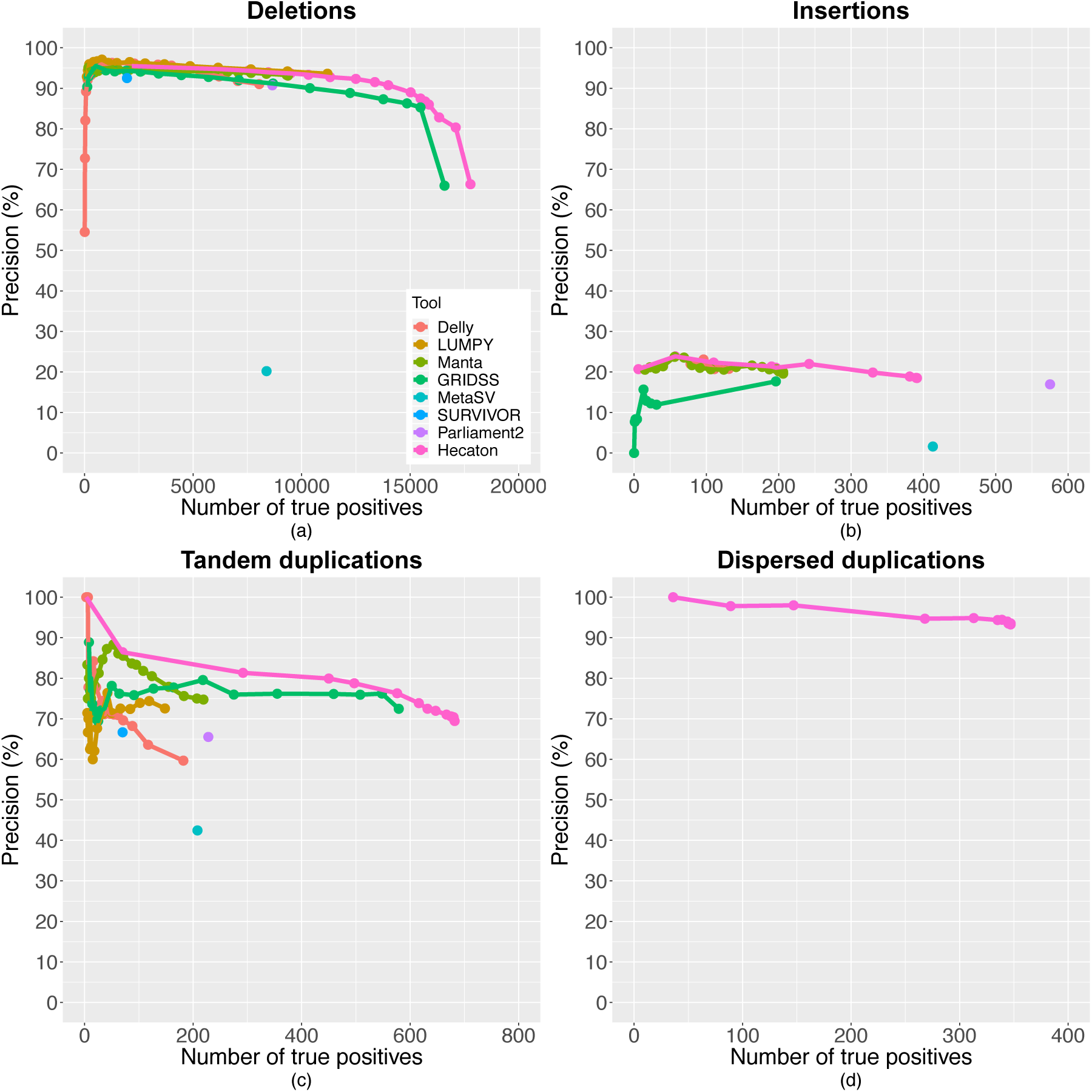
Performance of different CNV detection algorithms on short read data of the maize B73 accession. CNV calls were called relative to the maize Mo17 reference assembly. Precision-recall curves of Delly, LUMPY, Manta, and GRIDSS were constructed by varying the minimum number of discordantly aligned read pairs and/or split reads supporting each call in the set. The curve of Hecaton was produced by varying the threshold of the probabilistic score used to define calls as true positives. Performance is shown separately for deletions (a), insertions (b), tandem duplications (c), and dispersed duplications (d). Curves of LUMPY and SURVIVOR are missing for insertions, as these tools are unable to detect this type of CNV. Curves are missing for all tools besides Hecaton for dispersed duplications, as Hecaton is the only tool that can detect this type of CNV, owing to its post-processing stage. Hecaton generally improved recall and precision compared to Delly, LUMPY, Manta, and GRIDSS. Moreover, it significantly outperformed MetaSV, SURVIVOR, and Parliament2, three ensemble approaches applicable to plant data.

Consistent with previous benchmarks performed with long read data (Sedlazeck et al., 2018; De Coster et al., 2019), insertions remained difficult to reliably detect using short paired-end Illumina reads in all of our test cases, even after applying the filtering stage of Hecaton. We manually investigated alignments covering tens of false positive insertions in *A. thaliana* L*er* and discovered that they all resulted from alignments that were soft-clipped at the insertion site. These insertions were all reported by Hecaton to have an unknown size. With some of the insertions, the mates of the soft-clipped reads mapped to a different chromosome, indicating that some may be interchromosomal transpositions instead.

The idea of predictor combination has been already applied to improve detection of structural variants from exome sequencing data (Pounraja et al., 2019), detection of somatic single nucleotide variants (Ewing et al., 2015), inference of gene regulatory networks (Marbach et al., 2012), and predictive models of breast cancer prognosis (Margolin et al., 2013). Our results demonstrate that this concept can be used to improve CNV detection as well, contrasting a previous benchmark that combined structural variation methods through a majority vote (Lee et al., 2018). This suggests that the aggregation approach used by Hecaton is better suited to deal with the different treatment of each tool of specific types of CNV than a majority vote. A possible reason for this is that the random forest model employed by Hecaton can capture interactions between tools and CNV types to some extent during aggregation, while a majority vote assigns an equal weight to each tool. Such an approach does not work well if most tools are ill-suited to detect a specific type of CNV.

We demonstrated that Hecaton can be applied to any plant species, without needing to retrain the random forest model (Figure 5). The performance of Hecaton can be further improved, as it is relatively easy to extend it to include other CNV detection tools or to train new random forest models using additional plant data. Nevertheless, Hecaton has some limitations. First, it has limited sensitivity for dispersed duplications when applied to very low coverage (5x) data. Second, although Hecaton has no upper limit in terms of the size of CNV it can detect, we were not able to evaluate its performance on CNVs that were larger than 1 Mb, as such calls tended to be falsely validated by one of our validation methods, VaPoR (Additional file 1: Figure S6). Third, it is not able to detect insertions with both high sensitivity and precision, a limitation it shares with other CNV detection tools designed to work with short WGS data (Sedlazeck et al., 2018). Finally, we were not able to robustly assess the performance of Hecaton in polyploid plant species, as we could not find polyploid samples of which both short and long read data were publicly available. We expect that the performance of Hecaton on polyploids should be comparable to the performance reported in this work on diploids, if the polyploid sample does not show strong differences between haplotypes. To deal with additional biases found in more complex polyploid species, it may be worthwhile to obtain ground truth annotations in order to train random forest models specifically tailored to polyploids. Such annotations can be obtained by generating a small set of polyploid samples using both Illumina and PacBio sequencing platforms. Simulated data could serve as ground truth data as well, but we were unable to generate simulated CNVs that accurately represent the distribution of CNVs in real scenarios, a problem encountered in previous work that benchmarked CNV detection tools (Kosugi et al., 2019).

### 3.3 Hecaton provides a scalable method to detect CNVs in plant species

Hecaton scales well to crop genomes when using conventional computational server resources. By making extensive use of parallelization, it processed samples of both domesticated and wild tomatoes in reasonable time, taking a minimum of 7 h and a maximum of 40 h to process a single sample (Table 1), when using Hecaton with 13 cores (Intel^®^ Xeon^®^ CPU E5-2670 v3 @ 2.30 GHz) on a Linux server (Ubuntu 16.04). For comparison, it would have taken a minimum of 67 h and a maximum of 200 h to run Hecaton on a single core (Table 1).

**Table 1:** Hecaton running time and memory use.

| Sample | CPU time<br>(h) | Real time<br>(h) | Peak resident<br>set size (Gb) |
| --- | --- | --- | --- |
| PI158760 | 119.5 | 13.1 | 27.8 |
| LA2706 | 67.8 | 7.3 | 23.9 |
| TR00003 | 96.2 | 11.1 | 24.3 |
| LA4451 | 70.8 | 7.3 | 23.6 |
| LA2157 | 196.1 | 32.5 | 26.6 |
| LA0716 | 172.3 | 30.3 | 24.9 |
| LYC4 | 189.1 | 39.2 | 25.7 |

Although we do not know the true CNVs present in these samples, we estimated that Hecaton attained lower precision in the tomato samples than in the *A. thaliana* and rice sets. We considered events to be likely false positives if they did not have a clear and uniformly lower (deletions) or higher (duplications) read depth compared to the rest of the chromosome its located on or to its flanking regions, or if they showed excessive (over 1000x) read coverage. Based on these criteria, we estimated that 30% of deletions, 20% of tandem duplications, and 80% of dispersed duplications in the wild accession LYC4 sample were false positives, based on inspection of a random sample of 20 CNVs of each type.

Additional filtering steps based on the median read depth of CNV calls and the presence of gaps in the regions flanking calls removed a significant number of CNVs from the callsets of both the domesticated and wild accessions (Table 2). However, they had little to no effect when applied to the callsets of *A. thaliana* Col-0–Cvi-0 and Suijing18 rice (Additional file 1: Figure S7). Therefore, Hecaton does not perform these steps by default. They are only meant to be used when working with samples that are distantly related to the reference genome or for which the reference genome assembly contains a significant number of gaps.

**Table 2:** Number of CNVs detected in tomato samples before and after filtering.

| Sample | No filter | Read depth<br>filter | Read depth<br>and gap filter |
| --- | --- | --- | --- |
| PI158760 | 482 | 347 | 97 |
| LA2706 | 934 | 701 | 403 |
| TR00003 | 1487 | 1162 | 880 |
| LA4451 | 1789 | 1482 | 1095 |
| LA2157 | 7127 | 5372 | 4851 |
| LA0716 | 10064 | 8472 | 7910 |
| LYC4 | 11988 | 9950 | 9407 |

Hundreds to thousands of CNVs remained after performing additional filtering (Table 2), indicating that even a conservative, high confidence set of events (see Additional file 1: Figure S8 for an example of such an event) called by Hecaton can provide a sizable pool of genetic variation that can be further characterized. Given that the reference genome of tomato used in this work is based on a domesticated cultivar, an expectedly larger number of CNVs were found in the wild accessions (LA2157, LA0716, LYC4) than in the domesticated ones (PI158760, LA2706, TR00003, LA4451). Most of the CNVs between the tomato samples and the reference genome consisted of deletions (Table 3), following a similar trend as seen in the *A. thaliana* Col-0–Cvi-0, *A. thaliana* L*er*, and Suijing18 rice samples. No insertions were reported for any of the samples (Table 3). We expect that this result does not reflect the actual underlying biology, but was rather caused by the stringent cut-off used to filter calls. Most events overlapped with repetitive elements (Table 4), which is not unexpected as low-complexity regions are thought to be one of the prime mediators of the formation of CNV (Hastings et al., 2009). A smaller, but non-negligible, fraction of CNVs overlapped with genes and coding sequences (Table 4), providing potential leads for CNV events having a functional or phenotypic impact.

**Table 3:** Types of CNV detected in tomato samples after filtering.

| Sample | Deletions | Insertions | Tandem duplications | Dispersed duplications | Total |
| --- | --- | --- | --- | --- | --- |
| PI158760 | 82 | 0 | 15 | 0 | 97 |
| LA2706 | 295 | 0 | 106 | 2 | 403 |
| TR00003 | 777 | 0 | 103 | 0 | 880 |
| LA4451 | 901 | 0 | 193 | 1 | 1095 |
| LA2157 | 4261 | 0 | 561 | 29 | 4851 |
| LA0716 | 7168 | 0 | 712 | 30 | 7910 |
| LYC4 | 8508 | 0 | 878 | 21 | 9407 |

**Table 4:** Overlap of filtered CNVs with repeats, genes, and coding sequences (CDS) in tomato samples

| Sample | Sequence covered (Mb) | Repeats (% of total) | Genes (% of total) | CDS (% of total) |
| --- | --- | --- | --- | --- |
| PI158760 | 0.21 | 64.9 | 12.38 | 2.32 |
| LA2706 | 0.46 | 75.59 | 12.47 | 1.73 |
| TR00003 | 1.59 | 84.01 | 9.38 | 1.89 |
| LA4451 | 1.21 | 76.26 | 11.78 | 1.11 |
| LA2157 | 8.59 | 82.79 | 11.01 | 1.25 |
| LA0716 | 12.11 | 85.47 | 12.19 | 1.29 |
| LYC4 | 12.13 | 81.53 | 13.41 | 1.04 |

## 4 Conclusion

Hecaton is a computational workflow specifically designed to detect CNV from WGS data of plant genomes. It strongly improves upon the performance of current approaches primarily developed for human genomes, indicating that such tools cannot be directly applied to plant data without optimization. In contrast to several state-of-the-art tools, Hecaton correctly detects dispersed duplications. Moreover, the random forest model employed by Hecaton significantly improves upon current approaches in terms of sensitivity and precision. Finally, the running time and memory usage of Hecaton scales well to crop genomes, demonstrating its practical utility to plant research.

We anticipate that Hecaton is of immediate interest to both applied and fundamental research regarding the relationship between genotype and phenotype in plants. CNVs have been linked with several stress-resistant phenotypes in crop species (Mickelbart et al., 2015), including frost tolerance in wheat (Würschum et al., 2017), aluminum tolerance in maize (Maron et al., 2013), and boron tolerance in barley (Sutton et al., 2007). Hecaton can extensively query crop and wild germplasm for resistant loci, that can be characterized and eventually introgressed into elite cultivars. Besides its use in agricultural research, Hecaton may help to answer more fundamental questions regarding the role of CNV, as many characteristics regarding the role of CNV in plant adaptation are still relatively unknown (Gaut et al., 2018). Such characteristics include how stress affects the rate at which CNVs accumulate, the main molecular mechanisms that govern the creation of CNVs, and the evolutionary dynamics that determine whether CNVs become fixed within a population. Populations of wild and domesticated plant species (such as the 100 Tomato Genome project (Aflitos et al., 2014)) may provide excellent datasets to explore these topics, given that domestication is a fairly recent phenomenon involving artificial selection for a set of well-defined and well-characterized traits.

## Availability of data and code

To develop and evaluate Hecaton, we used publicly available short and long read datasets of an *A. thaliana* Col-0–Cvi-0 F1 hybrid (Chin et al., 2016), the *Japonica* rice variety Suijing18 (Nie et al., 2017), *A. thaliana* Landsberg *erecta* (L*er*) (Zapata et al., 2016), maize B73 (Jiao et al., 2017), and domesticated and wild tomato accessions (Aflitos et al., 2014). Additional file 2: Table S2 provides a full description of the datasets, including NCBI Short Read Archive accession numbers. The genomic assemblies and annotations of *A. thaliana* Col-0 (version TAIR10), *O. sativa Japonica* (version IRGSP-1.0) and maize Mo17 (Zm-Mo17-REFERENCE-CAU-1.0) can be respectively found at the European Nucleotide Archive under accession numbers GCA_000001735.2, GCA_001433935.1, and GCA_003185045.1. The genomic assembly (version SL3.0) and annotation (version ITAG3.20) of *S. lycopersicum* Heinz 1706 can be obtained from the FTP server of Sol Genomics Network (ftp://ftp.solgenomics.net/tomato_genome). The code repository of Hecaton can be found at git. wur.nl/bioinformatics/hecaton.

## Acknowledgements

We thank Carlos de Lannoy and Mehmet Akdel for testing the installation of Hecaton.

## Author contributions

R.W. implemented Hecaton, evaluated its performance and applied it to representative crop samples. All authors contributed to writing and reviewing the manuscript.

## Supplementary Material

### Additional file 1

**Figure S1:** Performance of different CNV detection algorithms on the test set of Col-0– Cvi-0 and Suijing18 CNV events called from 10x coverage data. **Figure S2:** Interpretating the appropriate type of CNV from a set of novel adjacencies. **Figure S3:** Recall of the post-processing step of Hecaton for dispersed duplications simulated at different allele dosages in tetraploid tomato genomes, before and after post-processing. **Figure S4:** Performance of Hecaton on the test set containing Col-0–Cvi-0 and Suijing18 CNV events called from 10x coverage data, using random forest models trained on CNVs detected at different levels of sequencing coverage. **Figure S5:** Performance of different CNV detection algorithms on *A. thaliana* L*er* data having 10x coverage. **Figure S6:** Percentage of false positive events in simulated data that were incorrectly labeled as a true positive by VaPoR. **Figure S7:** Effect of filtering CNV calls based on read depth and presence of gaps in their flanking regions in the test set of Col-0–Cvi-0 and Suijing18 generated from 10x coverage data. **Figure S8:** Example of a high-confidence CNV called by Hecaton. **Figure S9:** Percentage of true events in simulated data incorrectly labeled as false positives by VaPoR. **Figure S10:** Percentage of true events in simulated data incorrectly labeled as false positives by Sniffles, in a set of 10 simulated versions of *S. lycopersicum* with distinct sets of CNVs. **Figure S11:** Percentage of false positive events in simulated data incorrectly labeled as a true positive by Sniffles, computed in a set of 10 simulated versions of *S. lycopersicum* with distinct sets of CNVs. **Figure S12:** Density plots of normalized median read depths of CNV events called in domesticated and wild tomato samples. **Figure S13:** Density plot of the fraction of N’s in the 400 bp flanking regions of CNV events called in domesticated and wild tomato samples. (PDF 354.8 kb)

### Additional file 2

**Table S1:** Features used for the random forest model. **Table S2:** Description of the used datasets. **Table S3:** Number of events called from B73 data that could not be validated by VaPoR. **Table S4:** Number of events simulated per size interval. (PDF 52.3 kb)

### Additional file 3

**Supplementary Methods:** Evaluating the performance of CNV detection tools using simulated data. Evaluating the performance of CNV detection tools using real data. Applying Hecaton to tomato data. (PDF 118.0 kb)

