## Additional file 1 for "Hecaton: reliably detecting copy number variation in plant genomes using short read sequencing data"

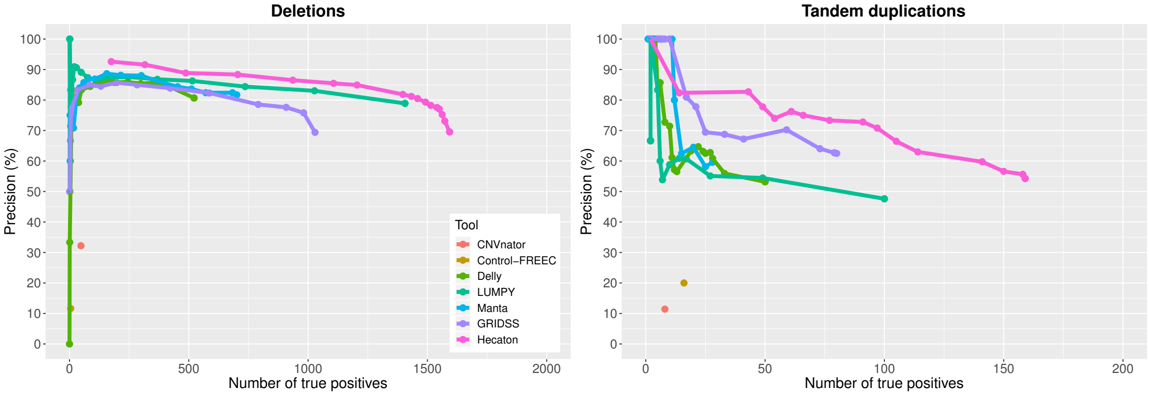


(a) (b)

Figure S1: **Performance of different CNV detection algorithms on the test set of Col-0–Cvi-0 and Suijing18 CNV events called from 10x coverage data.** Precision-recall curves of Delly, GRIDSS, LUMPY, and Manta were constructed by varying the minimum number of discordantly aligned read pairs and/or split reads supporting each call in the set. The curve of Hecaton was produced by varying the threshold of the probabilistic score used to define calls as true positives. Performance is shown separately for deletions and tandem duplications. CNVnator and Control-FREEC performed significantly worse than the other tools.

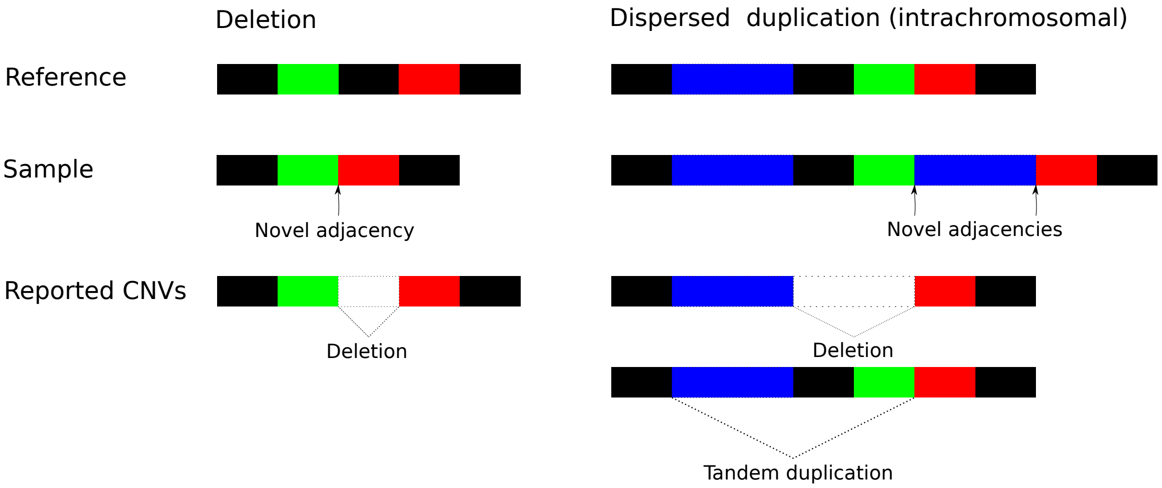


Figure S2: **Interpretating the appropriate type of CNV from a set of novel adjacencies.** A deletion introduces a single novel adjacency in the sample genome between the green and red segments. Delly, LUMPY, and Manta correctly interpret this adjacency as a deletion relative to the reference genome. Interchromosomal dispersed duplications introduce two novel adjacencies in the sample genome: one between the green and blue segment and one between the blue and red segment. Delly, LUMPY, and Manta interpret each of these adjacencies in isolation. As a result, they incorrectly interpret such adjacencies as overlapping pairs of false positive deletions and tandem duplications.

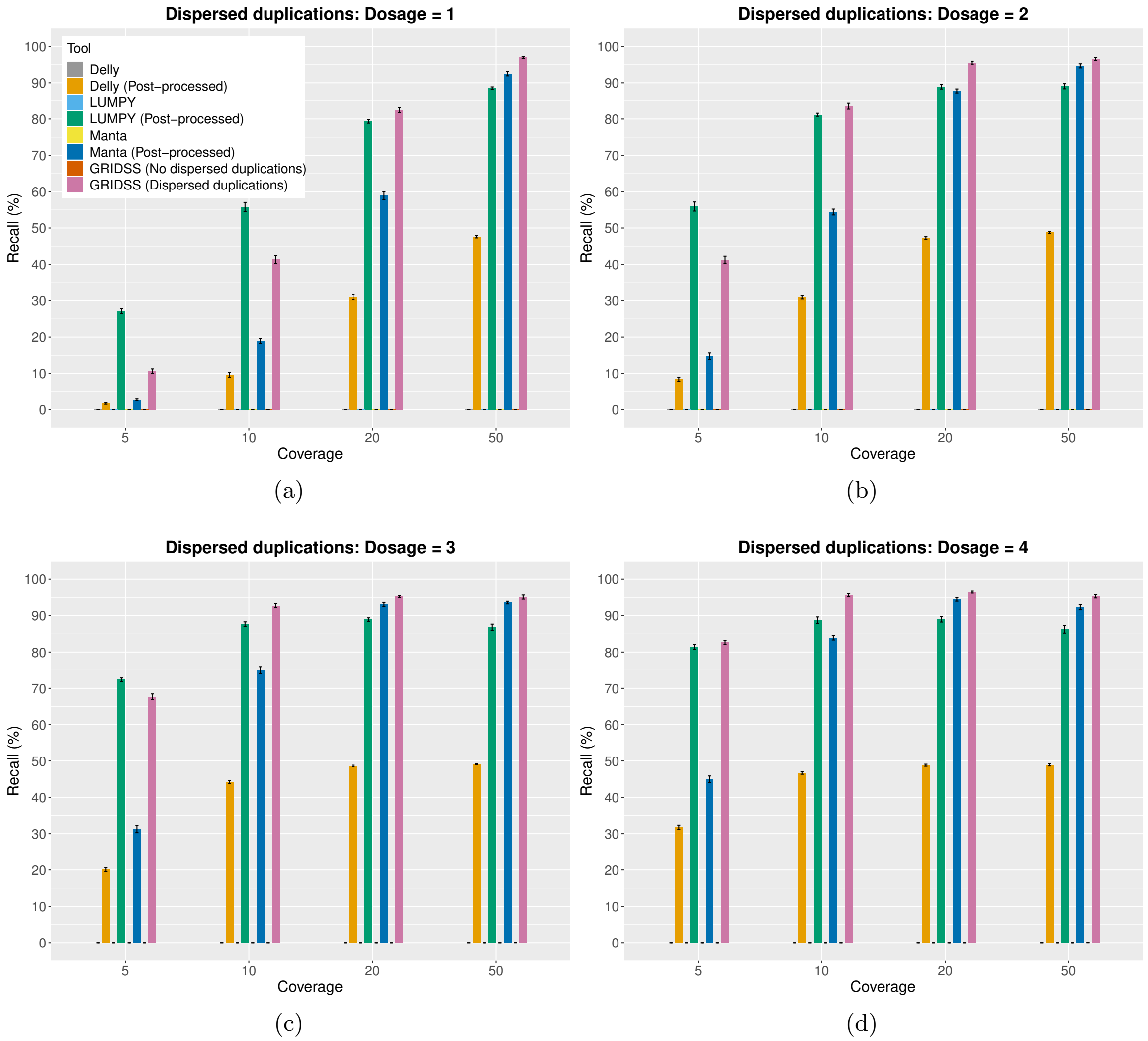


Figure S3: **Recall of the post-processing step of Hecaton for dispersed duplications simulated at different allele dosages in tetraploid tomato genomes, before and after post-processing.** Error bars depict the standard error of the mean. Results of GRIDSS were generated by processing adjacencies in isolation (no dispersed duplications) or by processing them in clusters (dispersed duplications). Recall is given for dispersed duplications simulated at dosage 1, 2, 3, and 4. As the coverage of variant alleles decreases with dosage, the post-processing step detected a lower fraction of the dispersed duplications simulated at low dosage compared to those simulated at high dosage.

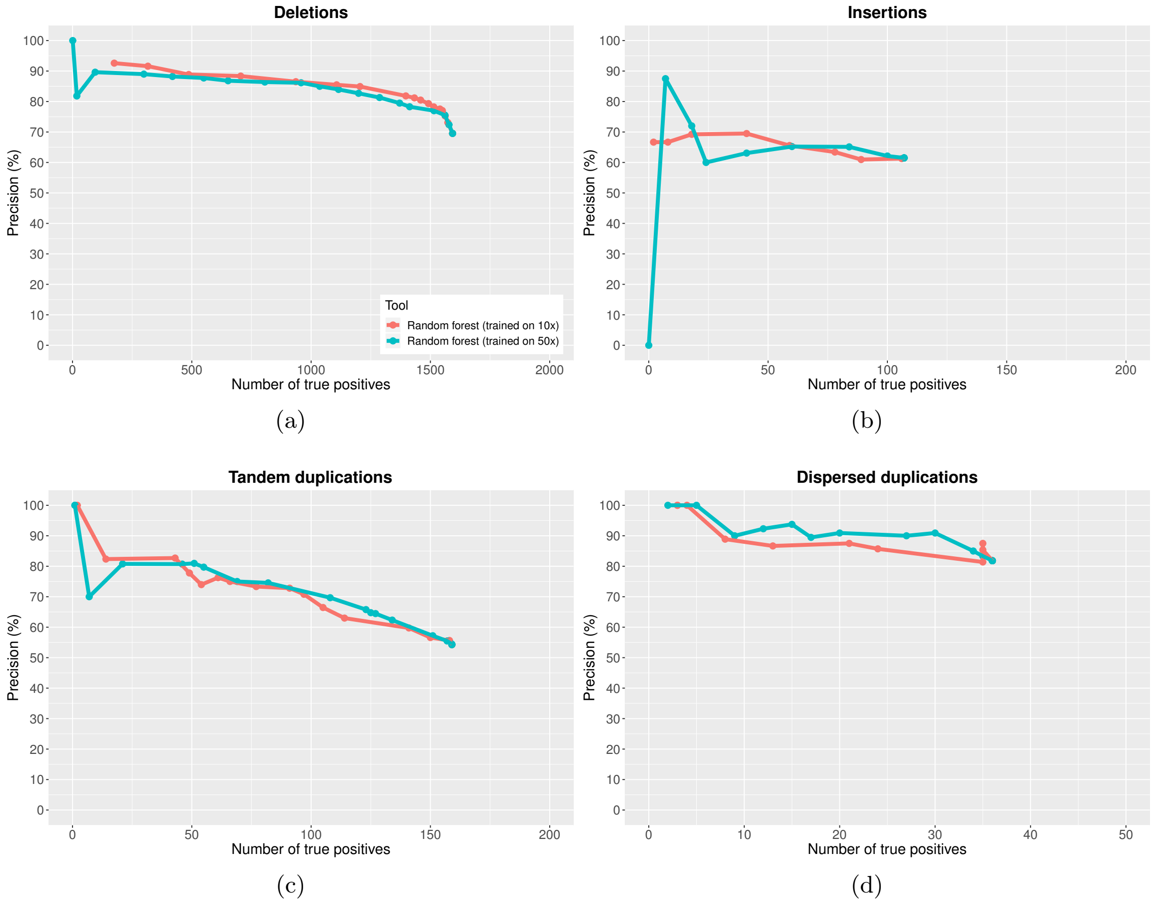


Figure S4: **Performance of Hecaton on the test set containing Col-0–Cvi-0 and Suijing18 CNV events called from 10x coverage data, using random forest models trained on CNVs detected at different levels of sequencing coverage.** The curves of both models were produced by varying the threshold of the probabilistic score used by Hecaton to define calls as true positives. Performance is shown separately for deletions (a), insertions (b), tandem duplications (c), and dispersed duplications (d). The two models perform about equally well.

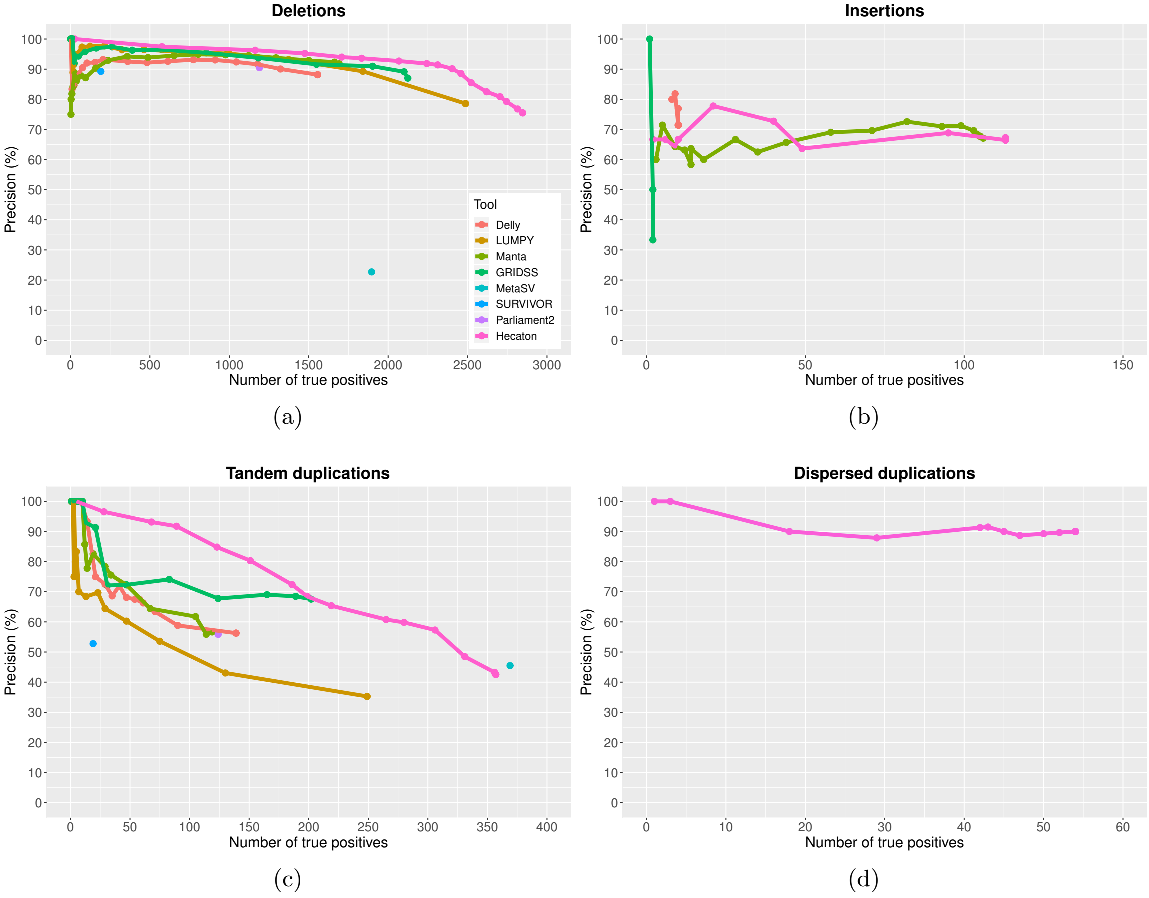


Figure S5: **Performance of different CNV detection algorithms on *A. thaliana* L*er* data at 10x coverage.** Precision-recall curves of Delly, LUMPY, Manta, and GRIDSS were constructed by varying the minimum number of discordantly aligned read pairs and/or split reads supporting each call in the set. The curve of Hecaton was produced by varying the threshold of the probabilistic score used to define calls as true positives. Performance is shown separately for deletions (a), insertions (b), tandem duplications (c), and dispersed duplications (d). The curve of LUMPY is missing for insertions, as this tools is unable to detect this type of CNV. Curves are missing for all tools besides Hecaton for dispersed duplications, as Hecaton is the only tool that can detect this type of CNV, owing to its post-processing stage. Hecaton generally improved recall and precision compared to Delly, LUMPY, Manta, and GRIDSS.

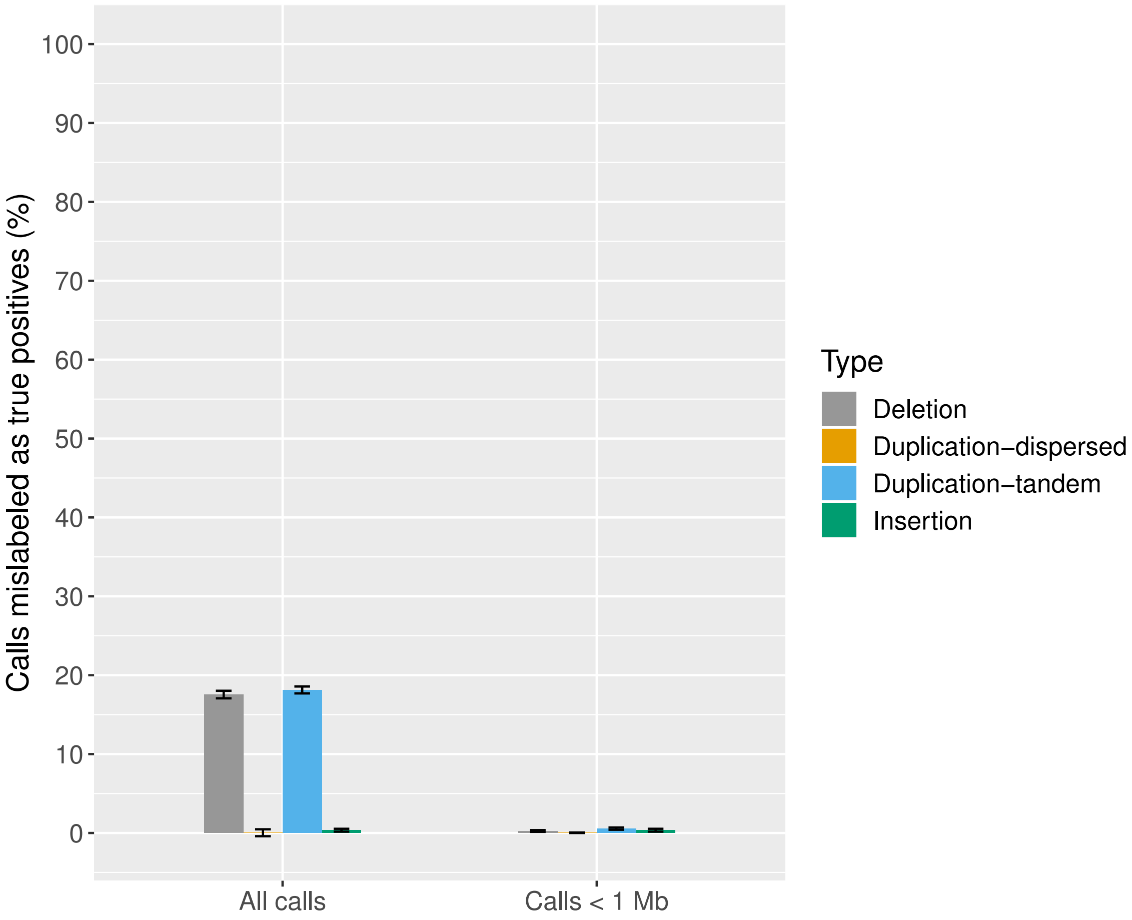


Figure S6: **Percentage of false positive events in simulated data that were incorrectly labeled as a true positive by VaPoR.** We computed the average percentage of false positive events incorrectly considered as true positives by VaPoR in a set of 10 simulated versions of *S. lycopersicum* with distinct sets of CNVs that were called by Manta and post-processed by Hecaton. The error bars depict the standard error of the mean. Most of the incorrectly labeled false positives were larger than 1 Mb.

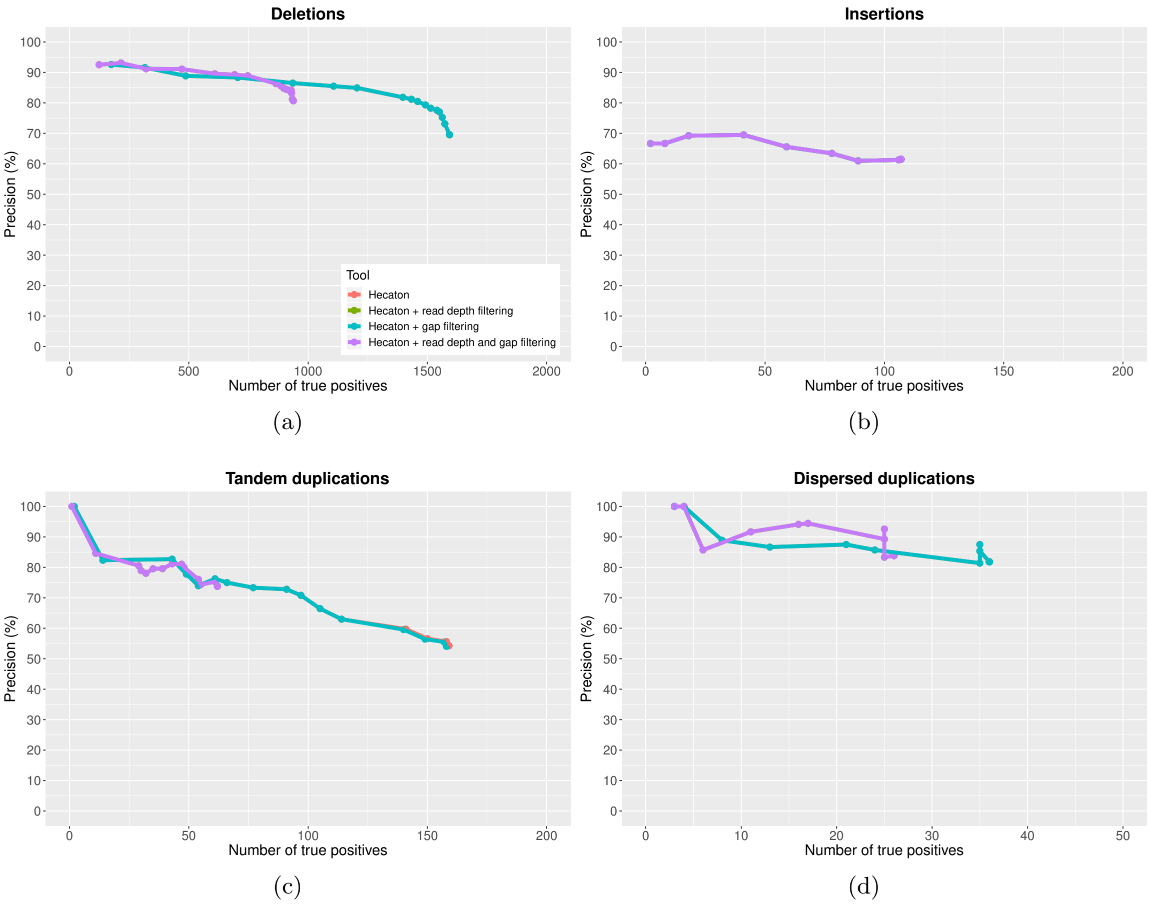


Figure S7: **Effect of filtering CNV calls based on read depth and presence of gaps in their flanking regions in the test set of Col-0–Cvi-0 and Suijing18 generated from 10x coverage data.** Curves were produced by varying the threshold of the probabilistic score used by Hecaton to define calls as true positives. Performance is shown separately for deletions, insertions, tandem duplications, and dispersed duplications. The curves did not change after gap filtering, due to the high quality of the used *A. thaliana* and rice reference genomes. Read depth filtering generally resulted in lower sensitivity, without significantly improving precision.

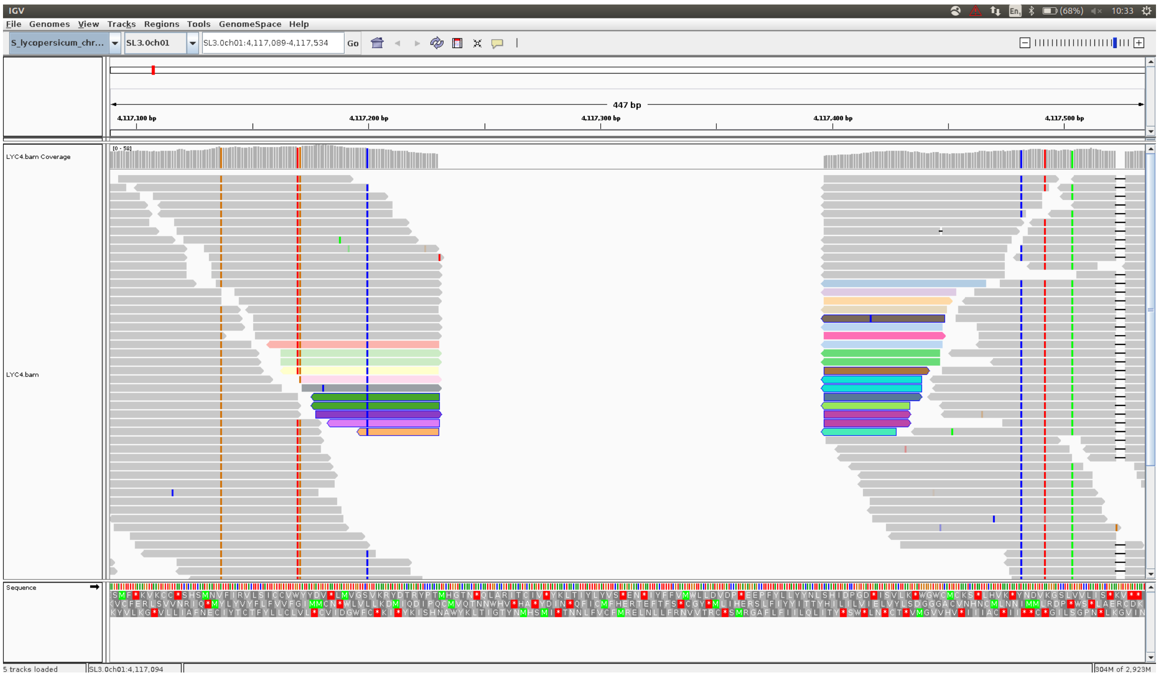


Figure S8: **Example of a high-confidence CNV called by Hecaton.** Hecaton called a deletion (region 4117226-4117397 of chromosome 1) in the wild tomato accession LYC4 relative to the *S. lycopersicum* Heinz 1706 reference genome (assembly version SL3.0). Alignments of LYC4 reads surrounding the deletion are visualized here using Integrative Genome Viewer (IGV). The deleted region (depicted in the middle of the figure) has a clear, uniformly lower read depth compared to that of its flanks, indicating that the deletion call is likely to be correct.

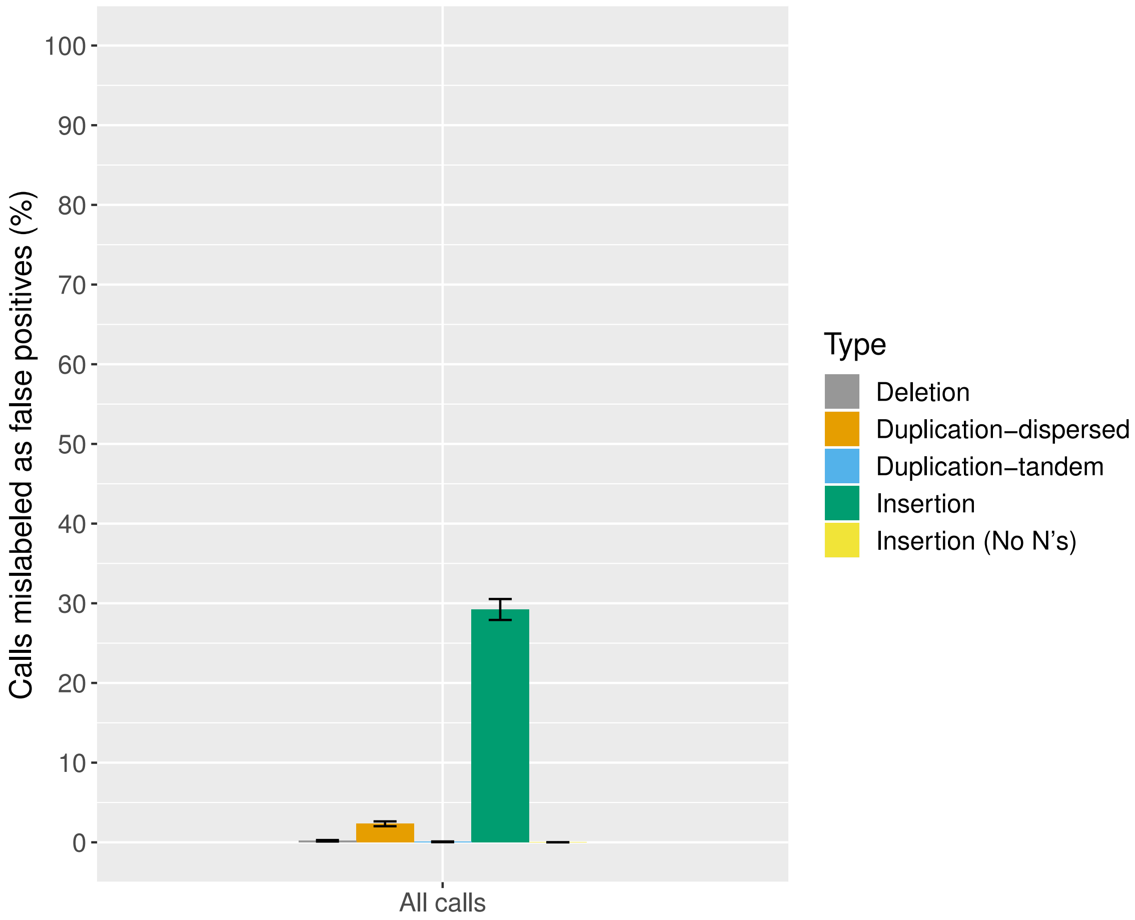


Figure S9: **Percentage of true events in simulated data incorrectly labeled as false positives by VaPoR.** We computed the average percentage of true events that were incorrectly seen as false positives by VaPoR in a set of 10 simulated versions of *S. lycopersicum* with distinct sets of CNVs that were called by Manta and post-processed by Hecaton. The error bars depict the standard error of the mean. Dispersed duplications were treated as insertions to allow them to be validated. VaPoR correctly labeled the large majority of events as true positive calls.

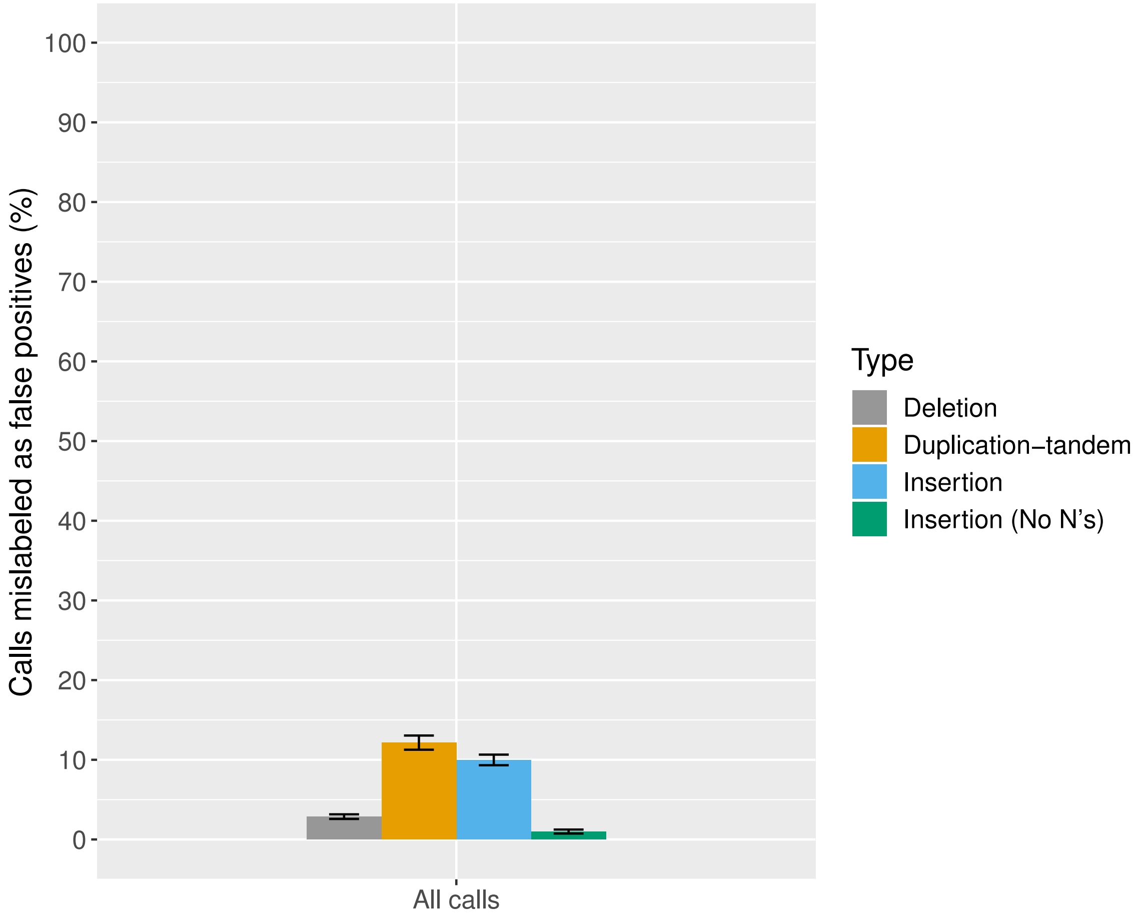


Figure S10: **Percentage of true events in simulated data incorrectly labeled as false positives by Sniffles, in a set of 10 simulated versions of *S. lycopersicum* with distinct sets of CNVs.** The error bars depict the standard error of the mean. Sniffles mislabeled only a small percentage of true positive insertions as false positives, of which most consisted of calls with unknown size.

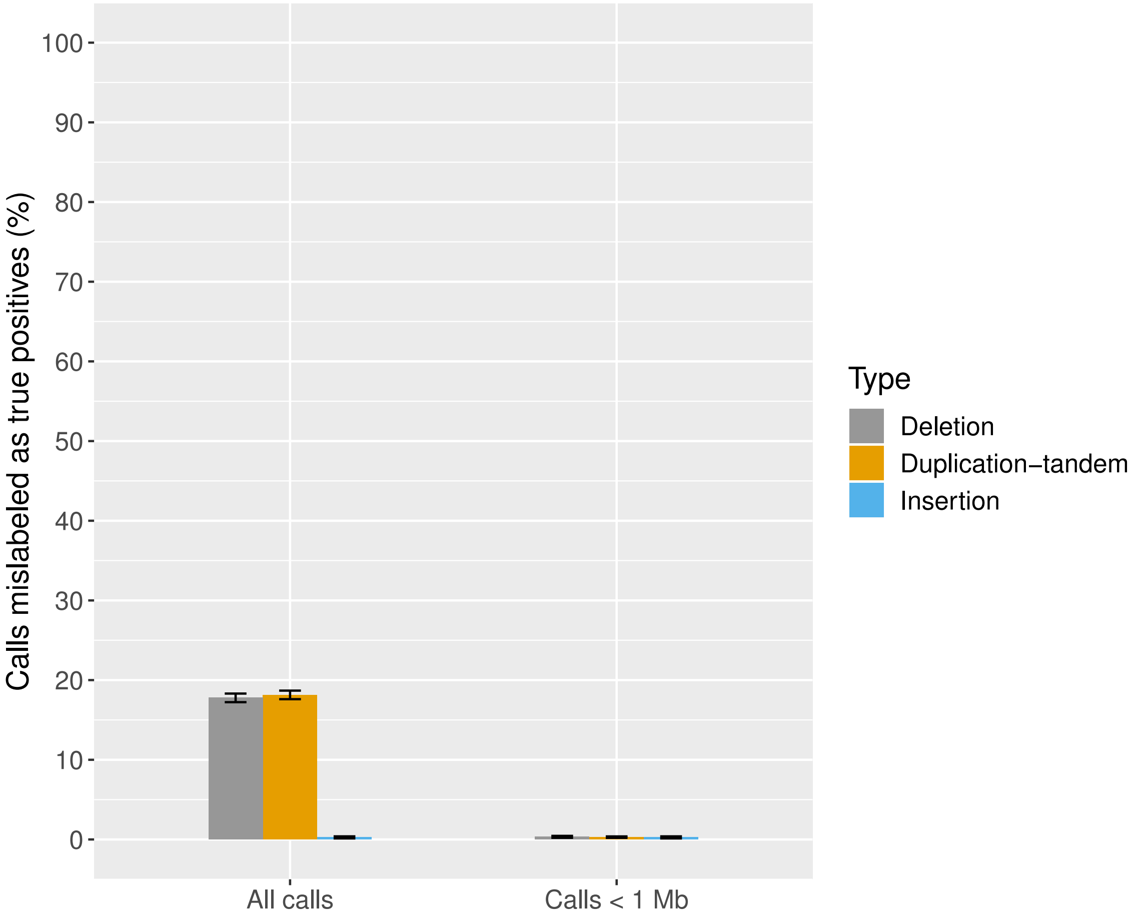


Figure S11: **Percentage of false positive events in simulated data incorrectly labeled as a true positive by Sniffles, computed in a set of 10 simulated versions of *S. lycopersicum* with distinct sets of CNVs.** The error bars depict the standard error of the mean. Sniffles correctly labeled all simulated true positive insertions, indicating that it accurately validates this type of event.

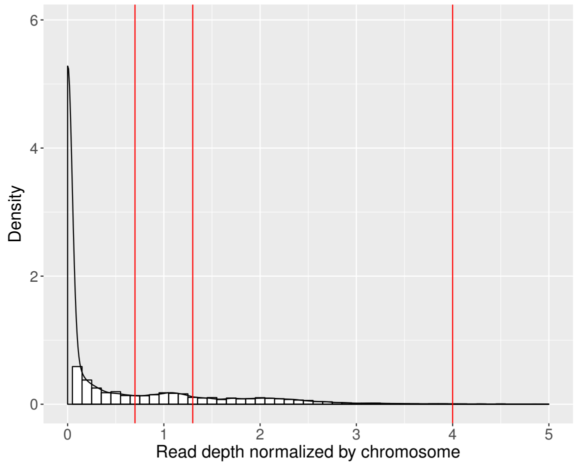


(a)

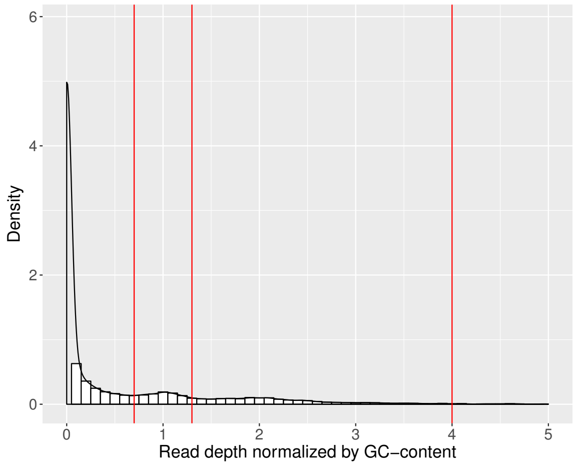


(b)

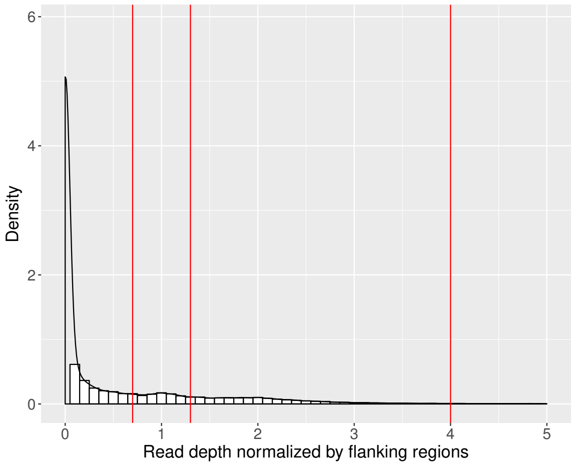


(c)

Figure S12: **Density plots of normalized median read depths of CNV events called in domesticated and wild tomato samples.** Densities are shown for read depths normalized by the depth of the chromosome the event is located on, regions in the genome with similar GC-content, and the 1000 bp regions flanking it. Red lines indicate the default cutoffs at which CNV events are filtered. All three plots show a small peak between 0.7 and 1.3. We considered CNVs having a median read depth value within any of the three peaks to be false positives.

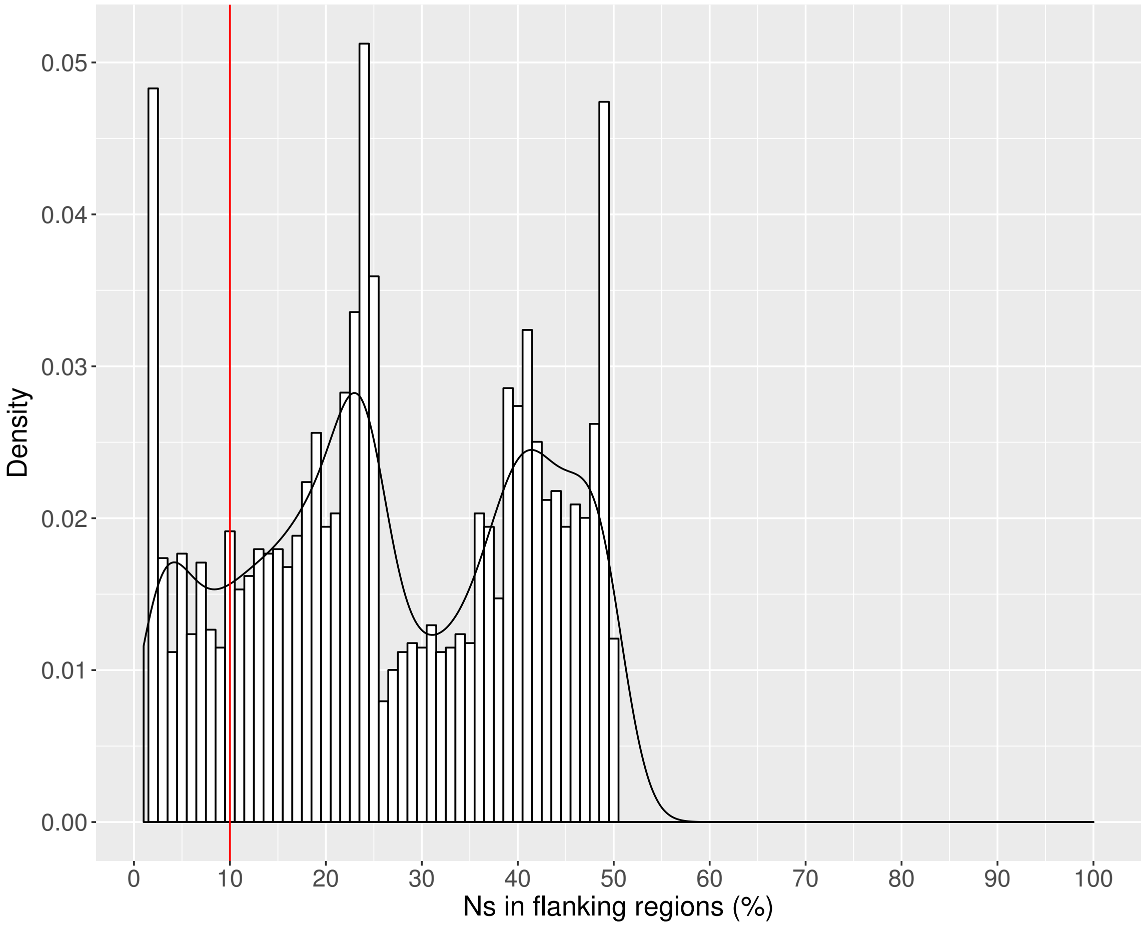


Figure S13: **Density plot of the fraction of N's in the 400 bp flanking regions of CNV events called in domesticated and wild tomato samples.** The plot was created after filtering events based on read depth. The frequency bar at 0% is omitted for clarity, as it is much larger than the frequency bars at all of the other values. The red line indicates the default cutoff at which CNV events are filtered. We expected that the presence of gaps correlates with the location of false positives, but that there is no relationship between the presence of gaps and the location of true positives. Therefore, we assumed that any peaks that appear in the distribution plot should be mainly caused by the presence of false positive CNVs. Based on this plot, we considered all CNVs of which at least 10 % of the flanking regions consisted of gaps to be likely false positives.
