## Additional file 2 for "Hecaton: reliably detecting copy number variation in plant genomes using short read sequencing data"

Table S1: **Features used for the random forest model**

| Feature | Type | Description |
|---|---|---|
| READ_PAIRS | Integer | Number of supporting discordantly aligned reads pairs |
| SPLIT_READS | Integer | Number of supporting split reads |
| SIZE | Integer | Size of the event in bp |
| DELLY | Binary | 1 if event is supported by Delly, 0 otherwise |
| GRIDSS | Binary | 1 if event is supported by GRIDSS, 0 otherwise |
| LUMPY | Binary | 1 if event is supported by LUMPY, 0 otherwise |
| MANTA | Binary | 1 if event is supported by Manta, 0 otherwise |
| DEL | Binary | 1 if event is a deletion, 0 otherwise |
| INS | Binary | 1 if event is an insertion, 0 otherwise |
| TANDUP | Binary | 1 if event is a tandem duplication, 0 otherwise |
| DISDUP | Binary | 1 if event is a dispersed duplication, 0 otherwise |

Table S2: **Description of the used datasets**

| Sample | Platform | Accession number | Avg. read length | Number of bases |
|---|---|---|---|---|
| *A. thaliana* Col-0–Cvi-0 | Illumina | SRX1865253 | 2 × 249 bp | 8.1 Gb |
| *A. thaliana* Col-0–Cvi-0 | PacBio | SRX1715706 | 5.6 kb | 15.7 Gb |
| *O. sativa* Suijing18 | Illumina | SRR5880534 | 2 × 150 bp | 23.6 Gb |
| *O. sativa* Suijing18 | PacBio | SRR5877285 | 8.5 kb | 26.4 Gb |
| *A. thaliana* L*er* | Illumina | SRR3166543 | 2 × 100 bp | 20.8 Gb |
| *A. thaliana* L*er* | PacBio | SRX533607 | 4.8 kb | 36.1 Gb |
| *Z. mays* ssp. *mays* L. B73 | Illumina | SRR2960981 | 2 × 250 bp | 209.8 Gb |
| *Z. mays* ssp. *mays* L. B73 | PacBio | SRX1472849 | 8.1 kb | 279.3 Gb |
| *S. lycopersicum* PI158760 | Illumina | ERR418074 | 2 × 100 bp | 35.9 Gb |
| *S. lycopersicum* LA2706 | Illumina | ERR418039 | 2 × 100 bp | 31.4 Gb |
| *S. lycopersicum* TR00003 | Illumina | ERR418051 | 2 × 100 bp | 33.3 Gb |
| *S. lycopersicum* LA4451 | Illumina | ERR418065 | 2 × 100 bp | 29.1 Gb |
| *S. arcanum* LA2157 | Illumina | ERR418092 | 2 × 100 bp | 32.0 Gb |
| *S. habrochaites* LYC4 | Illumina | ERR418105 | 2 × 100 bp | 31.8 Gb |
| *S. pennellii* LA0716 | Illumina | ERR418107 | 2 × 100 bp | 24.9 Gb |

Table S3: **Number of events called from B73 data that could not be validated by VaPoR**

| Tool | Number of events | % of total |
|---|---|---|
| Delly | 18074 | 44.41 |
| GRIDSS | 43006 | 48.45 |
| LUMPY | 7635 | 24.08 |
| Manta | 6957 | 23.74 |
| MetaSV | 35234 | 28.56 |
| Survivor | 2658 | 32.22 |
| Parliament2 | 12869 | 34.12 |
| Hecaton | 54322 | 51.13 |

Table S4: **Number of events simulated per size interval**

| Size interval | Number of events |
|---|---|
| 100-200 bp | 10 |
| 200-500 bp | 10 |
| 500-1000 bp | 10 |
| 1-2 kb | 10 |
| 2-5 kb | 10 |
| 5-10 kb | 5 |
| 10-20 kb | 5 |
| 20-50 kb | 3 |
| 50-100 kb | 3 |
