## Additional file 3 for "Hecaton: reliably detecting copy number variation in plant genomes using short read sequencing data"

### Supplementary Methods

#### Evaluating the performance of CNV detection tools using simulated data

##### Data generation

To evaluate CNV detection algorithms in a controlled setting, we simulated a large number of next-generation sequencing datasets based on rearranged versions of the *S. lycopersicum* Heinz 1706 reference genome (assembly version SL3.0) of tomato (The Tomato Genome Consortium, 2012). Although simulations do not contain all of the variation found in real sequencing data, they provide a way to obtain a lower bound of the performance of CNV detection callers in real scenarios. In order to perform many simulations in a reasonable amount of time, we generated datasets from the two largest chromosomes (1 and 9) only. Gap regions were excluded from simulations.

The following parameters were varied during simulations:

- **Ploidy** Both a diploid and a tetraploid version of chromosome 1 and 9 were simulated with a SNP rate of 0.02 using the haplogenerator.py script of Haplosim (git commit id 7bbc639f) (Motazedi et al., 2017), reflecting the varying levels of ploidy found among plant genomes. The distances between SNPs were modeled using a Poisson distribution with $\lambda$ chosen according to the chosen SNP rate. For simplicity, all SNPs were biallelic. In the diploid genome, 50% of SNPs were homozygous relative to the reference sequence and 50% were heterozygous. In the tetraploid genome, monoplex, duplex, triplex and quadruplex SNPs were simulated in equal proportions, assigning 25% of SNPs to each category.

- **Variant size and type** Deletions, insertions, single-copy tandem duplications, intrachromosomal dispersed duplications, and interchromosomal dispersed duplications were drawn from different size intervals (Additional file 2: Table S4). Events were randomly distributed to different genomic locations using bedtools (Quinlan and Hall, 2010) (version 2.27.1) shuffle. After being assigned a genomic location, events were simulated using the R-package RSVsim (Bartenhagen and Dugas, 2013) (version 1.14.0). The novel sequences inserted during insertion events were sampled from a randomly generated genomic sequence of 100 Mb. To capture the effect of variant dosage, a fraction of events were simulated in only a subset of the haplotype sequences. In diploid genomes, one half of the variants were simulated in one of the haplotypes (heterozygous variants) and the other half in both (homozygous variants). In tetraploid genomes, events were equally distributed over the following dosages: monoplex, duplex, triplex, and quadraplex. To prevent biasing the results towards one particular choice of genomic locations, 10 sets of variants were generated for each ploidy level.

- **Coverage** Next-generation sequencing datasets were simulated from each of the rearranged genomes using ART (Huang et al., 2011) (version 2.5.8). This tool mimics the sequencing protocol of Illumina sequencing platforms to generate a set of reads with realistic technical biases. To cover both low and high coverages, genomes were "sequenced" *in silico* at 5x, 10x, 20x and 50x coverage, using the paired-end HiSeq 2500 protocol with the read length set at 150 bp, the mean insert size at 400 bp and the standard deviation of the insert size at 40 bp.

Eighty different datasets were simulated in total, all having a different combination of ploidy level, variant sizes, genomic locations, and coverage. Each dataset was aligned to the reference sequence of *S. lycopersicum* Heinz 1706 chromosomes 1 and 9 using the Speedseq pipeline (Chiang et al., 2015) (version 0.1.2) with default parameters, similar to the alignment step in the calling stage of Hecaton.

##### Benchmarking tools

We applied Delly, LUMPY, Manta, and GRIDSS to simulated data and converted their output to a common representation using the post-processing step of Hecaton. To assess whether the post-processing step improved the detection of dispersed duplications, it was run with and without correction of erroneously represented CNVs. Delly, LUMPY, Manta, and GRIDSS were all applied to the simulated datasets using minimum filtering parameters, in order to minimize variation caused by the choice of parameters for each different tool.

We assessed the performance of tools using two different metrics: recall and precision. Recall is defined as the number of true positives found by a tool divided by the number of simulated events. A detected variant was labeled as a true positive if it had more than 50% reciprocal overlap with a simulated variant of the same type. Otherwise, it was labeled as a false positive. Precision is defined as the number of detected true positives divided by the total number of detected variants.

#### Evaluating the performance of CNV detection tools using real data

##### Validation tools

To determine whether CNV events called in real data were true or false positives, we employed two tools that validate calls detected from short Illumina paired-end read data by using long PacBio reads. We evaluated deletions, tandem duplications and dispersed duplications using VaPoR (Zhao et al., 2017), which uses alignments of PacBio reads to validate deletion, duplication, and insertion events called from Illumina data. However, VaPoR is unable to validate insertions of which the size of the inserted sequence could not be exactly determined by the CNV detection algorithm. As this is the case for a large fraction of insertions detected from Illumina reads, we evaluated insertions using a different tool, Sniffles (Sedlazeck et al., 2018), that directly calls structural variants from alignments of PacBio reads. We kept CNVs discovered by Sniffles if they were supported by at least 5 PacBio reads.

Both VaPoR and Sniffles require alignments of PacBio data to the same reference genome that was used to produce the calls based on Illumina data. We used NGMLR (Sedlazeck et al., 2018), a mapper optimized for detecting structural variation, to generate such alignments. In the case of *A. thaliana* L*er*, the Illumina and PacBio data were generated from two different L*er* genotypes, respectively L*er*-1 and L*er*-0, as there was no publicly available PacBio data for the Illumina sample used in this study. We do not expect that using these two different genotypes led to any problems regarding validation, as only a marginal amount of structural variation was found between them (Zapata et al., 2016). Deletions, tandem duplications, and dispersed duplications that were assigned a score of at least 0.15 by VaPoR were labeled as true positives, as recommended by the developers of VaPoR (Zhao et al., 2017). It should be noted that VaPoR only validates the insertion site of a dispersed duplication and not whether the location of the template sequence of the duplication was called correctly. Insertions were labeled as true positives if they were located within 50 bp of an insertion detected by Sniffles.

To determine the rate at which VaPoR and Sniffles correctly label false and true positives, we used them to validate events called by Manta from simulated 10x coverage Illumina datasets of *S. lycopersicum*, using PacBio reads simulated from the same genome with PBSIM (Ono et al., 2012) (40x coverage, Data type: CLR). VaPoR tends to falsely validate calls larger than 1 Mb (Additional file 1: Figure S6). To prevent such calls from influencing our results, we excluded them from our evaluation. VaPoR was able to correctly validate the large majority of true positive events (Additional file 1: Figure S9), as only a small percentage of true dispersed duplications were incorrectly labeled as false positives. Sniffles mislabeled only a small percentage (10 %) of true positive insertions as false positives, most of which consisted of calls with unknown size (Additional file 1: Figure S10). It did not label any false positive insertions as true positive (Additional file 1: Figure S11). Based on these findings, we concluded that both VaPoR and Sniffles accurately validate events called from Illumina data and are appropriate tools for generating a ground truth callset that can be used to optimize and evaluate Hecaton.

##### Benchmarking tools

Two metrics were used to evaluate the performance of tools: the number of true positives (calls that were validated by VaPoR or Sniffles) and the precision. The number of true positives serves to approximate the recall of a particular tool when applied to real datasets, as we do not know the true number of CNVs in real data. Precision is defined as the number of true positives divided by the total number of calls that could be validated by VaPoR or Sniffles. VaPoR is unable to validate calls located in regions too rich in repeats. We excluded such unvalidated calls from our evaluation.

We evaluated the performance of random forest models and individual tools on the held-out test sets of Col-0–Cvi-0 and Suijing18 and on data of *A. thaliana* L*er* and maize B73. To evaluate performance on low coverage data, which is often encountered in plant studies, all datasets were subsampled to 10x coverage using seqtk (Li, 2012). The output of individual tools can contain calls that are highly similar to each other, in some cases differing only by a single base pair. We consider such calls to be duplicates in practice. To prevent such duplicates from influencing results, we merged calls using the same criteria as those used in the merging step of Hecaton before evaluating each tool. Precision-recall curves of individual tools were constructed by varying the minimum number of discordantly read pairs and split reads supporting each call in the output. Curves for random forest models were produced by varying the threshold of the probabilistic score used to filter calls from the final output.

For the Col-0–Cvi-0 and Suijing18 data, we applied the random forest models to the features of the held-out calls and compared the results to the labels that were assigned to these calls by VaPoR and Sniffles. In order to determine whether the sequencing coverage used during model training has a significant effect on performance, we evaluated two different random forest models: one trained on 10x coverage data and one trained on 50x coverage data. The precision and number of true positives of individual tools were computed by only taking into account calls located on chromosomes present in the held-out set. In the case of the *A. thaliana* L*er* and maize B73 samples, we evaluated both the models and the individual callers using all chromosomes, as these samples are independent from the ones used to train the random forest models. For the maize B73 sample, a large fraction of calls could not be validated by VaPoR due to the highly repetitive nature of the used reference assembly (Additional File 2: Table S3). Therefore, we only included calls of B73 that overlapped for at least 50% of their length with genes or the 5000 bp interval upstream or downstream of genes, using bedtools (version 2.27.1) slop and bedtools intersect. We believe that this benchmark is still a representative measure of performance, as downstream analysis of CNVs detected by short reads generally focuses on genic, non-repetitive regions.

To compare the performance of Hecaton to that of other ensemble methods used to detect CNVs, we used the same datasets to benchmark three methods that aggregate the results of different tools: MetaSV (Mohiyuddin et al., 2015), SURVIVOR (Jeffares et al., 2017) and Parliament2 (Zarate et al., 2018). We would have liked to compare the performance of Hecaton to that of FusorSV (Becker et al., 2018) as well, but this particular ensemble method is unfortunately not applicable to plant data.

MetaSV (version 0.5.4) was evaluated by first running BreakDancer (version 1.4.5-unstable-66-4e44b43 (commit 4e44b43)), Pindel (0.2.5b9, 20160729), and CNVnator (v0.3.2) on alignments generated by the calling stage of Hecaton. The resulting callsets were integrated by running MetaSV with its soft-clip detection and local assembly features turned on. CNVnator was run using a window size of 200. Pindel and MetaSV require the mean and standard deviation of the insert size of the used datasets as parameters; these were computed with Picard CollectInsertSizeMetrics (version 2.9.2) using a sample of 1 million reads for each dataset. All other parameters of the tools were set to their default values.

SURVIVOR (version 1.0.5) was used to generate a consensus callset out of the output of Delly, LUMPY, and Manta. All three callers were run using default parameters. We ran SURVIVOR with the same parameters used by its developers in previous work (Jeffares et al., 2017; Sedlazeck et al., 2018). With these parameters, calls between the tools are collapsed into a single call if they are of the same type and if their start and end coordinates are within 1 kb of each other. Consensus calls were kept if they were supported by least two out of three tools.

We ran Parliament2 (version 0.1.9-13-g37d63065) with the versions of BreakDancer, Delly, LUMPY, and Manta that were included with its Docker image, merging calls using the –genotype parameter. The resulting calls were merged using the same criteria as those used in the merging step of Hecaton to prevent double counting calls that likely correspond to the same CNV event.

#### Applying Hecaton to tomato data

To demonstrate the utility of Hecaton, we applied it to short read data of both domesticated and wild tomato accessions (Additional file 2: Table S2). We chose to apply Hecaton to these samples for two reasons. As the size of the reference genome of tomato (828 Mb) is significantly larger than that of *A. thaliana* (119 Mb) and rice (374 Mb), applying Hecaton to tomato samples should indicate whether its run time and memory requirements scale well to crop data. Moreover, as copy number variation is thought to be involved in many of the phenotypic adaptations underlying the domestication of crops (Gaut et al., 2018), these datasets should provide an accurate expectation of the number of CNVs we should expect to find in crop samples.

##### Detecting and characterizing CNVs

To obtain a lower bound on the number of calls that we can expect to find in tomato samples, we aimed to find a conservative, high confidence set of CNVs. We detected CNVs in each sample relative to the *S. lycopersicum* Heinz 1706 reference genome (assembly version SL3.0) using Hecaton. We only kept CNVs which had a score of 0.75 in the output of each sample, as filtering calls using this threshold resulted in a precision of at least 80% for all four types of CNVs in the datasets of Col-0–Cvi-0, Suijing18, and L*er*. To get an indication of the biological relevance of detected CNVs, we computed the overlap of CNV events with annotated repeat and gene models (version ITAG3.2) using bedtools (version 2.27.1) intersect.

##### Benchmarking running time and memory

The running time and memory usage of Hecaton was determined by using the Nextflow command line options ”-with-report” and ”-with-trace”. These options report CPU time, wall clock time and the resident set size of each executed process. The three metrics are internally computed by the Nextflow framework using the unix commands “ps” and “date”.

##### Filtering CNVs based on read depth

To filter CNVs based on read depth, we computed the read depth of regions involved in CNV using duphold (Pedersen and Quinlan, 2019) (v0.1.1). This tool computes the median read depth of a CNV event and normalizes it by dividing it by three different values: the median read depth of the rest of the chromosome that the event is located on, the median read depth of regions in the genome with similar GC-content, and the median read depth of the 1000 bp regions flanking it. We expected that a true positive CNV event has normalized read depths that differ significantly from 1. Therefore, we filtered calls if any of the three normalized read depths had a value between 0.7 and 1.3. These thresholds were chosen based on distribution plots of the three normalized read depths of CNVs detected in domesticated and wild tomato accessions (Additional file 1: Figure S12). The plots showed a small peak between these values, which we assumed to be false positive CNV events. We additionally filtered calls of which any of the normalized read depths had a value higher than 4, indicating spurious coverage.

##### Filtering CNVs based on the presence of gaps in their flanking regions

To filter CNVs based on the presence of gaps in their flanking regions, we extracted the 200 bp regions flanking the 5' and 3' side of each breakpoint using bedtools (version 2.27.1) flank and bedtools getfasta. Next, we computed the fraction of Ns in these flanking regions using a custom Python script. Finally, we filtered calls if at least 10% of their flanking regions consisted of Ns. This threshold was chosen based on a distribution plot of the fraction of Ns in flanking regions of CNVs detected in domesticated and wild tomato accessions (Additional file 1: Figure S13). We expected that the presence of gaps correlates with the location of false positives, but that there is no relationship between the presence of gaps and the location of true positives. Therefore, we assumed that any peaks that appear in the distribution plot should be mainly caused by the presence of false positive CNVs. As such peaks were found to the right of the value 0.1 in the plot (Additional file 1: Figure S13), we chose to set the filtering threshold to 10%.
